# IL-1β-driven NF-κB transcription of ACE2 as a Mechanism of Macrophage Infection by SARS-CoV-2

**DOI:** 10.1101/2024.12.24.630260

**Authors:** Cadence Lee, Rachel Khan, Chris S. Mantsounga, Sheila Sharma, Julia Pierce, Elizabeth Amelotte, Celia A. Butler, Andrew Farinha, Crystal Parry, Olivya Caballero, Jeremi A. Morrison, Saketh Uppuluri, Jeffrey J. Whyte, Joshua L. Kennedy, Xuming Zhang, Gaurav Choudhary, Rachel M. Olson, Alan R. Morrison

## Abstract

Coronavirus disease 2019 (COVID-19), caused by infection with the enveloped RNA betacoronavirus, SARS-CoV-2, led to a global pandemic involving over 7 million deaths. Macrophage inflammatory responses impact COVID-19 severity; however, it is unclear whether macrophages are infected by SARS-CoV-2. We sought to identify mechanisms regulating macrophage expression of ACE2, the primary receptor for SARS-CoV-2, and to determine if macrophages are susceptible to productive infection. We developed a humanized *ACE2* (*hACE2*) mouse whereby *hACE2* cDNA was cloned into the mouse *ACE2* locus under control of the native promoter. We validated the susceptibility of *hACE2* mice to SARS-CoV-2 infection relative to wild-type mice and an established *K18-hACE2* model of acute fulminating disease. Intranasal exposure to SARS-CoV-2 led to pulmonary consolidations with cellular infiltrate, edema, and hemorrhage, consistent with pneumonia, yet unlike the *K18-hACE2* model, *hACE2* mice survived and maintained stable weight. Infected *hACE2* mice also exhibited a unique plasma chemokine, cytokine, and growth factor inflammatory signature relative to *K18-hACE2* mice. Infected *hACE2* mice demonstrated evidence of viral replication in infiltrating lung macrophages, and infection of macrophages in vitro revealed a transcriptional profile indicative of altered RNA and ribosomal processing machinery as well as activated cellular antiviral defense. Macrophage IL-1β-driven NF-κB transcription of ACE2 was an important mechanism of dynamic ACE2 upregulation, promoting macrophage susceptibility to infection. Experimental models of COVID-19 that make use of native hACE2 expression will allow for mechanistic insight into factors that can either promote host resilience or increase susceptibility to worsening severity of infection.

## INTRODUCTION

The new enveloped RNA betacoronavirus, severe acute respiratory syndrome coronavirus 2 (SARS-CoV-2), was first identified and brought to the world’s attention upon outbreak in the city of Wuhan, China, in December of 2019 (1–3). By March of 2020, the World Health Organization (WHO) declared coronavirus disease 2019 (COVID-19), the disease associated with the SARS-CoV-2 infection, a global pandemic, and over the ensuing 3 years, approximately 7 million deaths worldwide were directly attributed to this disease (4). It is estimated that about 80% of infections are asymptomatic or associated with mild upper respiratory symptoms like cough, headache, ageusia, anosmia, dyspnea, fatigue, and fever (1, 5–8). However, a subset of patients manifests moderate-to-severe clinical presentations ranging across the spectrum of pneumonia, acute respiratory distress syndrome (ARDS), septic shock with multi-organ failure, and death (9). As the virus has moved into an endemic cycle, about one in ten Americans has reported experiencing post-acute sequela of COVID-19 (PASC), referred to as “long COVID”, with about 27% of those affected noting significant limitations in their day-to-day activities (10). Several factors appear to influence the heterogeneity of presentation across the spectrum of the host, the environment, and the evolution of the virus (11). While the mechanistic determinants of the wide variability in clinical presentation remain unclear, it was evident early in the pandemic that age, age-associated comorbidities like cardiovascular disease, diabetes, and lung disease, as well as underlying genetic polymorphisms, correlated with increased risk of severe disease (12). Initial case-fatality reports for patients with coronary artery disease, diabetes mellitus, and hypertension were 10.5%, 7.3%, and 6.0%, respectively, and all were higher than the overall initial case-fatality of 2.3% (13). Current estimates now indicate that coronary artery disease is associated with a 12-fold increased risk of death and that myocardial infarction is present in 22% of patients requiring care in the intensive care unit (14).

Severe presentations of COVID-19 have been associated with a maladapted induction of an immune response to the infection analogous to ‘‘cytokine storm’’ or cytokine release syndrome (CRS), which defined an influenza-like syndrome occurring after systemic infections and immunotherapies (15–18). During the early stages of infection, host interferon activation and signaling were lower than expected and possibly insufficient to clear the virus, leading to a consequent enhanced inflammatory cytokine, chemokine, and growth factor activation (19, 20). While this damaging systemic elevation in cytokines has been associated with the host response to COVID-19, whether this is truly a cytokine storm remains controversial as the concentrations of cytokines like TNF-α, IL-6, and IL-8 do not appear to be as strong as those found in sepsis, non-COVID-19 ARDS, trauma, cardiac arrest, and CRS (21–24). Immune responses associated with COVID-19 exhibit dynamic and time-dependent alterations in the systemic levels of many cytokines, including IL-6, the kinetics of which are not fully understood, and thus, our mapping of this process is still incomplete (25).

Hyperinflammation associated with COVID-19 has been associated with increases in multiple cytokines, chemokines, and growth factors, including IL-1α, IL-1β, IL-6, IL-7, TNF*-*α, type I and II IFNs, CCL2, CCL3, and CXCL10 (25–27). Genetic findings, interaction maps of viral proteins with host factors, and high-resolution single-cell transcriptomics analyses have indicated that the host innate immune system, particularly the myeloid compartment, may play a role in determining disease severity and outcomes (28–31). In fact, accumulating data indicate that airway macrophages and recruited infiltrating monocyte-derived macrophages appear to be a key source of this hyperinflammatory cytokine expression implicated in more severe presentations of COVID-19 (30, 32–34).

Whether phagocytosis of cellular material and virion particles derived from infected cells promotes maladaptive hyperinflammation or whether innate immune cells are actual targets of productive infection with SARS-CoV-2 remains controversial. Early immunostaining of post-mortem tissue from patients who had died from COVID-19 identified co-staining of CD169^+^ macrophages with SARS-CoV-2 nucleocapsid protein (35). CD68^+^ macrophages from kidneys of patients with COVID-19 and acute kidney injury also co-stained positive for viral nucleocapsid (36). Some have called into question the expression of angiotensin-converting enzyme 2 (ACE2), the primary receptor for SARS-CoV-2, in macrophage populations, and in vitro studies indicate infection of monocyte-derived macrophages and dendritic cells (DCs) with SARS-CoV-2 is abortive and does not support virus replication (20, 31, 34, 37, 38). More recent studies involving lung tissue culture models suggest that activated macrophages are infected by the virus, promoting an inflammatory cytokine positive feedback cycle (39).

SARS-CoV-2 is closely related to other coronaviruses such as SARS-CoV-1 and the Middle East respiratory syndrome coronavirus (MERS-CoV), which have also caused severe respiratory diseases and are known to be transmitted from animals to humans (40). Early studies of SARS-CoV-1 revealed ACE2 to be a major cell entry receptor for this family of viruses. The mechanism of cell entry involved engagement of ACE2 by the coronavirus spike protein after employing transmembrane protease serine 2 (TMPRSS2) for spike protein priming (41, 42). The interface between viral spike protein and ACE2 has been structurally characterized, and the efficiency of ACE2 usage was found to be a key determinant of SARS-CoV-1 transmissibility (43). SARS-CoV-2 shares 79.6% sequence identity to SARS-CoV-1, and the SARS-CoV-2 spike protein shares 76% amino acid identity with that of SARS-CoV-1 (44, 45). Several amino acid substitutions in the receptor-binding domain between viral spike protein and the surface of ACE2 support stronger binding affinity of SARS-CoV-2 relative to SARS-CoV-1 (46–48). Moreover, SARS-CoV-2 has adapted to make use of multiple proteases in addition to TMPRSS2, including cathepsin L, cathepsin B, trypsin, factor X, elastase, and furin for spike protein priming to facilitate ACE2 binding and cell entry. Due to key structural differences resulting from amino acid variations, mouse ACE2 protein does not support efficient SARS-CoV-1 or SARS-CoV-2 binding, presenting a challenge to the use of mice as genetic tools in the study of experimental COVID-19 (49, 50). Infection of wild-type strains of mice, including BALB/c and C57BL/6, with SARS-CoV-2 revealed no overt clinical manifestations, weight loss, or mortality, and viral presence appeared contained to the lungs and was rapidly cleared (51). The mouse and human *ACE2* genes are located on the X chromosome in the region neighboring the pseudo-autosomal region and are remarkably similar in structure and encoded protein domains (52, 53). Both human and mouse *ACE2* promoter sequences include two evolutionarily conserved regions, referred to as the distal and proximal promoter regions upstream of the *ACE2* translational start codon (54, 55).

Dynamic upregulation of ACE2 expression, particularly increased expression in response to inflammatory signaling or aging, may be a mechanism whereby macrophages become susceptible to infection in a manner comparable to airway epithelial cells (56, 57). We sought to understand the regulation of ACE2 expression in macrophages in response to inflammation and whether macrophages are infected by SARS-CoV-2. We generated a humanized *ACE2* (*hACE2*) mouse model whereby human *ACE2* cDNA was cloned into the native mouse *ACE2* gene locus under regulation of the mouse promoter. We assessed the expression of hACE2 in macrophages derived from *hACE2* mice and compared response to infection with SARS-CoV-2 with wild-type (C57BL/6) control mice and the widely used *K18-hACE2* [B6.Cg-Tg(K18-ACE2)2Prlmn/J] mouse, which is a transgenic mouse overexpressing hACE2 under control of the keratin18 (K18) promoter and is susceptible to infection by both SARS-CoV-1 and SARS-CoV-2 (58, 59).

## RESULTS

### Human ACE2 mRNA and protein are expressed in the lungs of *hACE2* mice

To develop an animal model of human ACE2 expression under regulation of the native mouse promoter, we generated a mouse strain in which the human *ACE2* cDNA was inserted into exon 2 of the native mouse gene locus, using a clustered regularly interspaced short palindromic repeat and its associated protein 9 (CRISPR/Cas9) strategy (**Fig. S1**). The insertion site in exon 2 was selected based on the availability of CRISPR guides to ensure a low off-target effect and conservation profile. Insertion of human *ACE2* cDNA was intended to result in expression of human *ACE2* mRNA in place of the mouse *ACE2* mRNA under control of the native mouse promoter. To validate the exchange of human in place of mouse *ACE2* mRNA expression, primer-specific RT-qPCR was carried out on RNA isolated from lung tissue homogenates using C57BL/6J and *K18-hACE2* mice as controls (**Fig. 1A-C**). Lung tissue from the *hACE2* mice demonstrated a complete loss of mouse *ACE2* mRNA relative to both the C57BL/6J and the *K18-hACE2* mice, both of which have intact mouse *ACE2* alleles. C57BL/6J mice did not express human *ACE2* mRNA as expected, whereas the *K18-hACE2* and *hACE2* mice demonstrated expression of human *ACE2* mRNA. *K18-hACE2* mice, consistent with their design as an overexpression system, expressed about 3-fold higher levels of human *ACE2* mRNA relative to the *hACE2* mice. To confirm that mRNA expression adequately reflects the expression of protein, we quantified human ACE2 protein in the airways of *hACE2* mice, using immunofluorescence microscopy of the lung fields with an antibody that we validated by immunoblot assay to be highly specific to human ACE2 relative to mouse ACE2 (**Fig S2**). Immunofluorescence microscopy demonstrated hACE2 protein to be associated with airways marked by α-tubulin in both the *hACE2* mice and the *K18-hACE2* mice but not in the C57BL/6J control mice (**Fig. 1D, E**).

**Figure 1.**
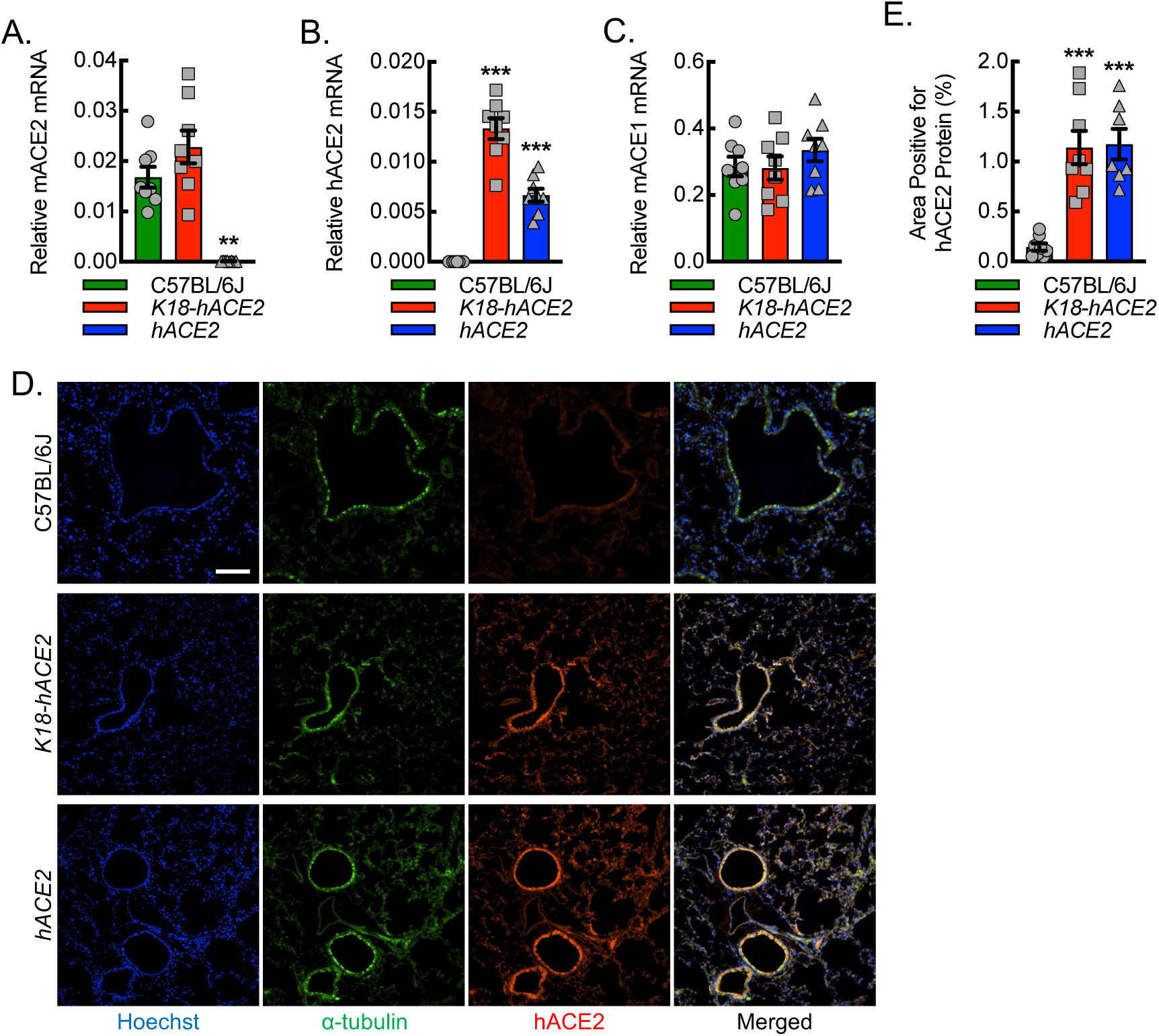
Validation of human ACE2 mRNA and protein expression in the lungs of *hACE2* mice. Primer-specific RT-qPCR for relative mRNA expression of mouse ACE2 (mACE2) (A), human ACE2 (hACE2) (B), and mouse ACE1 (mACE1) (C) from lung tissue homogenates of wild-type control (C57BL/6J) mice, K-18 promoter-driven hACE2 overexpression (*K18-hACE2*) mice, and humanized ACE2 (*hACE2*) mice (**, P<0.0005; ***, P<0.0001; compared to C57BL/6J by ANOVA; n=8, 4 male and 4 female mice). (D) Representative photomicrographs of hACE2 (red) expression in lung airways marked for alpha-tubulin (green) with corresponding quantitative data (E). Hoechst, blue. Bar, 200 microns. Data, mean ± SD.

### Infection of *hACE2* mice with SARS-CoV-2 is associated with increased survival and maintenance of weight relative to infection of *K18-hACE2* mice

Having validated the expression of human *ACE2* mRNA and protein in the lung tissue of *hACE2* mice, we next sought to determine whether *hACE2* mice were susceptible to infection by SARS-CoV-2, using the USA_WA1/2020 variant strain of the virus. C57BL/6J, *K18-hACE2,* and *hACE2* mice, 12 weeks of age, were intranasally exposed to either mock infection with vehicle control or 1000 PFU/dose of SARS-CoV-2 and then followed over 14 days for survival and changes in body weight (**Fig. 2**). At this dose of SARS-CoV-2 exposure, *K18-hACE2* mice exhibit severe infection with 87.5% mortality between days 6-8 post-infection along with approximately 20-25% weight loss, based on a dose-response curve and consistent with prior literature (**Fig. S3**) (58–62). Unlike the *K18-hACE2* mice, *hACE2* mice did not exhibit mortality or significant weight loss in response to the same SARS-CoV-2 exposure, but rather, they exhibited 100% survival and weight profiles that were more comparable to mock controls or infected C57BL/6J mice, demonstrating stability and a trend toward continued weight gain from baseline.

**Figure 2.**
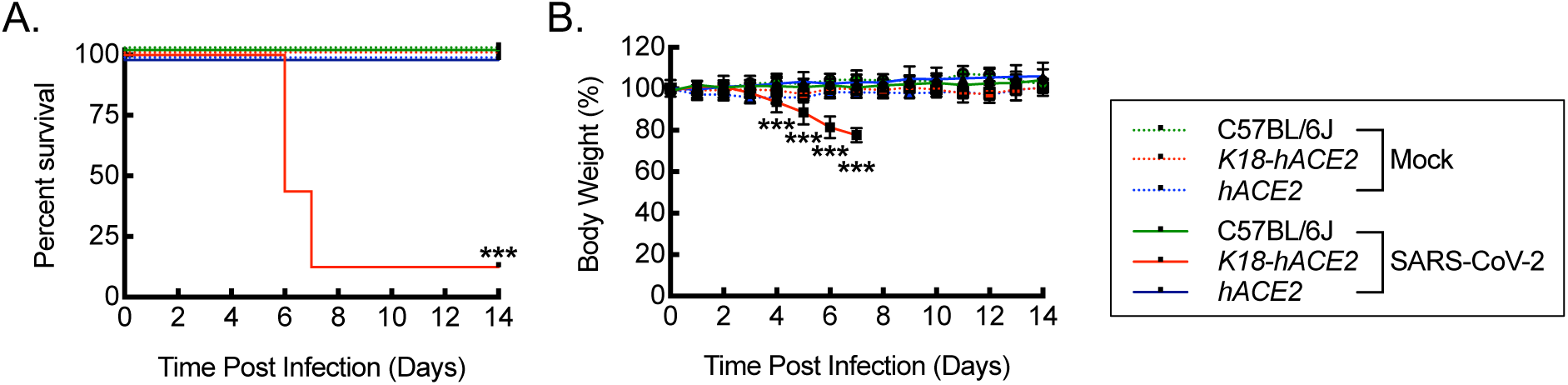
Infection of *hACE2* mice with SARS-CoV-2 is associated with increased survival and maintenance of weight relative to infection of *K18-hACE2* mice. (A) Kaplan-Meier plot for survival post intranasal infection of C57BL/6J, *K18-hACE2*, and *hACE2* mice with 1000 PFU/dose of SARS-CoV-2 or mock infection with vehicle control (***, P<0.0001 by log rank compared to mock or C57BL/6J controls; n=16, 8 male and 8 female, for SARS-CoV-2 infections; n=8, 4 male and 4 female, for vehicle control). (B) Percent body weight relative to pre-infection weight over time after SARS-CoV-2 infection carried out as in (A) (***, P<0.0001 compared to mock or C57BL/6J controls using ANOVA; n=16, 8 male and 8 female, for SARS-CoV-2 infections; n=8, 4 male and 4 female, for vehicle controls). Data, mean ± SD.

### *hACE2* mice are susceptible to SARS-CoV-2 infection

Because *hACE2* mice did not exhibit mortality or weight loss, we next sought to determine whether *hACE2* mice are susceptible to SARS-CoV-2 infection by quantitating viral RNA. C57BL/6J, *K18-hACE2,* and *hACE2* mice were subjected to mock or SARS-CoV-2 infection, and then RNA was isolated from lung tissue homogenates on day 6 post-infection. RT-qPCR was performed for the detection of viral RNA, using primers specific for open reading frame 1 (*ORF1*), spike (*S*), and nucleocapsid (*N*) RNA (**Fig. 3**). Mock infection served to establish a negative baseline. Both C57BL/6J and *hACE2* mice demonstrated significant and comparable expression of *ORF1*, *S*, and *N* RNA relative to baseline, with the *hACE2* mice trending modestly higher than in the C57BL/6J mice. Moreover, *K18-hACE2* mice demonstrated a further 2-3 log increase in *ORF1*, *S*, and *N* RNA detection over the C57BL/6J and *hACE2* mice, supporting a higher viral burden associated with the epithelial overexpression of hACE2 in the *K18-hACE2* mice.

**Figure 3.**
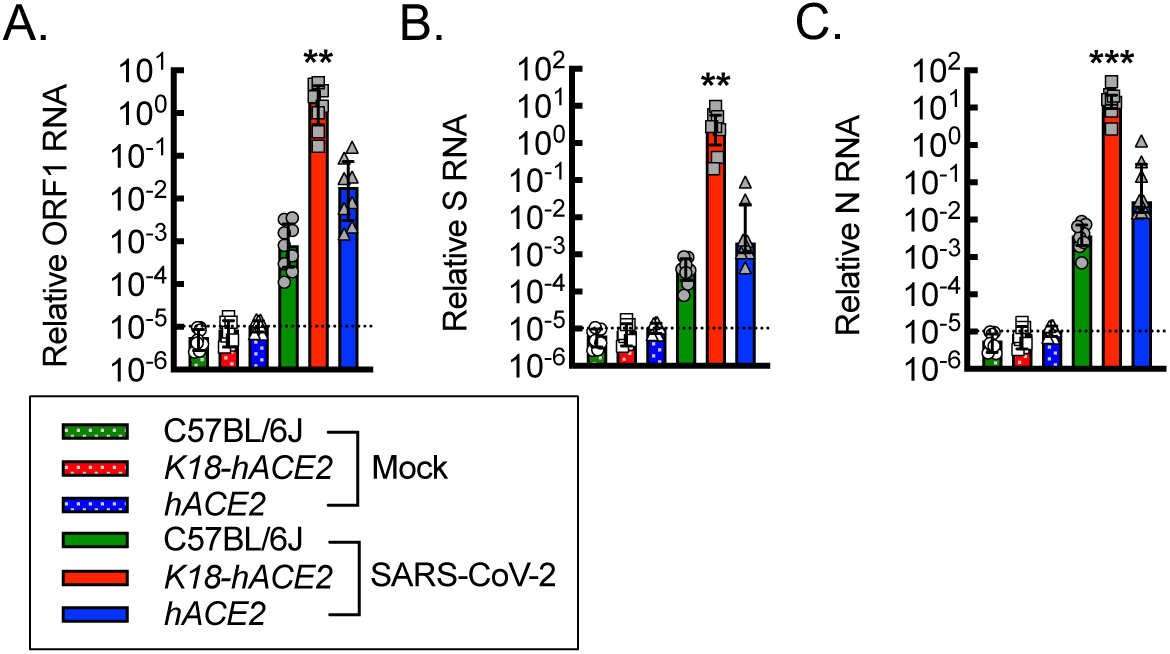
*hACE2* mice are susceptible to SARS-CoV-2 infection. RT-qPCR for relative SARS-CoV-2 *ORF1* (A), *S* (B), and *N* (C) RNA from lung tissue homogenates isolated from mice on day 6 post intranasal infection with vehicle control (Mock) or 1000 PFU/dose of SARS-CoV-2 (**, P<0.002; ***, P≤0.0005; relative to all others using ANOVA; n=8, 4 male and 4 female). Dotted line, average negative baseline on mock controls. Data, mean ± SD.

### Lung tissue from *hACE2* mice infected with SARS-CoV-2 exhibits comparable measures of pneumonia relative to *K18-hACE2* mice

To assess the severity of pulmonary infection associated with SARS-CoV-2 in the *hACE2* mice, we carried out quantitative histopathologic analyses of lung tissue specimens from mock and SARS-CoV-2-infected mice (**Fig. 4**). C57BL/6J, *K18-hACE2,* and *hACE2* mice, 12 weeks of age, were subjected to intranasal mock infection with vehicle control or 1000 PFU/dose of SARS-CoV-2, and lung tissue was harvested on day 6 post-exposure. Hematoxylin and eosin staining of lung tissue sections revealed areas of pulmonary consolidation that exhibited evidence of cellular infiltrate, edema, and alveolar hemorrhage, in both the *K18-hACE2* and *hACE2* mice infected with SARS-CoV-2 relative to C57BL/6J mice or mock vehicle control infections. Quantitation of nuclei per high-powered field as a marker of inflammatory cell infiltrate revealed a 2-fold increase in nuclei in the *K18-hACE2* and *hACE2* mice relative to C57BL/6J mice or mock-infected animals, supporting increased cellularity. Quantitation of airspace as an inverse marker for edema and tissue consolidation revealed a 25% reduction in airspace in both the *K18-hACE2* and *hACE2* mice relative to C57BL/6J mice or mock-infected animals. Masson’s trichrome staining allowed assessment of acute collagen deposition as part of the inflammatory response associated with SARS-CoV-2 infection. Quantitation of percent lung area staining blue for collagen revealed a 1.5-2-fold increase in the *K18-hACE2* and *hACE2* mice relative to C57BL/6J mice or mock-infected animals. Finally, immunohistochemistry staining for SARS-CoV-2 viral nucleocapsid protein revealed a 2-3-fold increase in percent area staining positive for viral nucleocapsid in the *K18-hACE2* and *hACE2* mice relative to C57BL/6J mice and an over ten-fold increase in signal relative background staining in the mock infections, supporting increased lung viral burden in both the *K18-hACE2* and *hACE2* mice.

**Figure 4.**
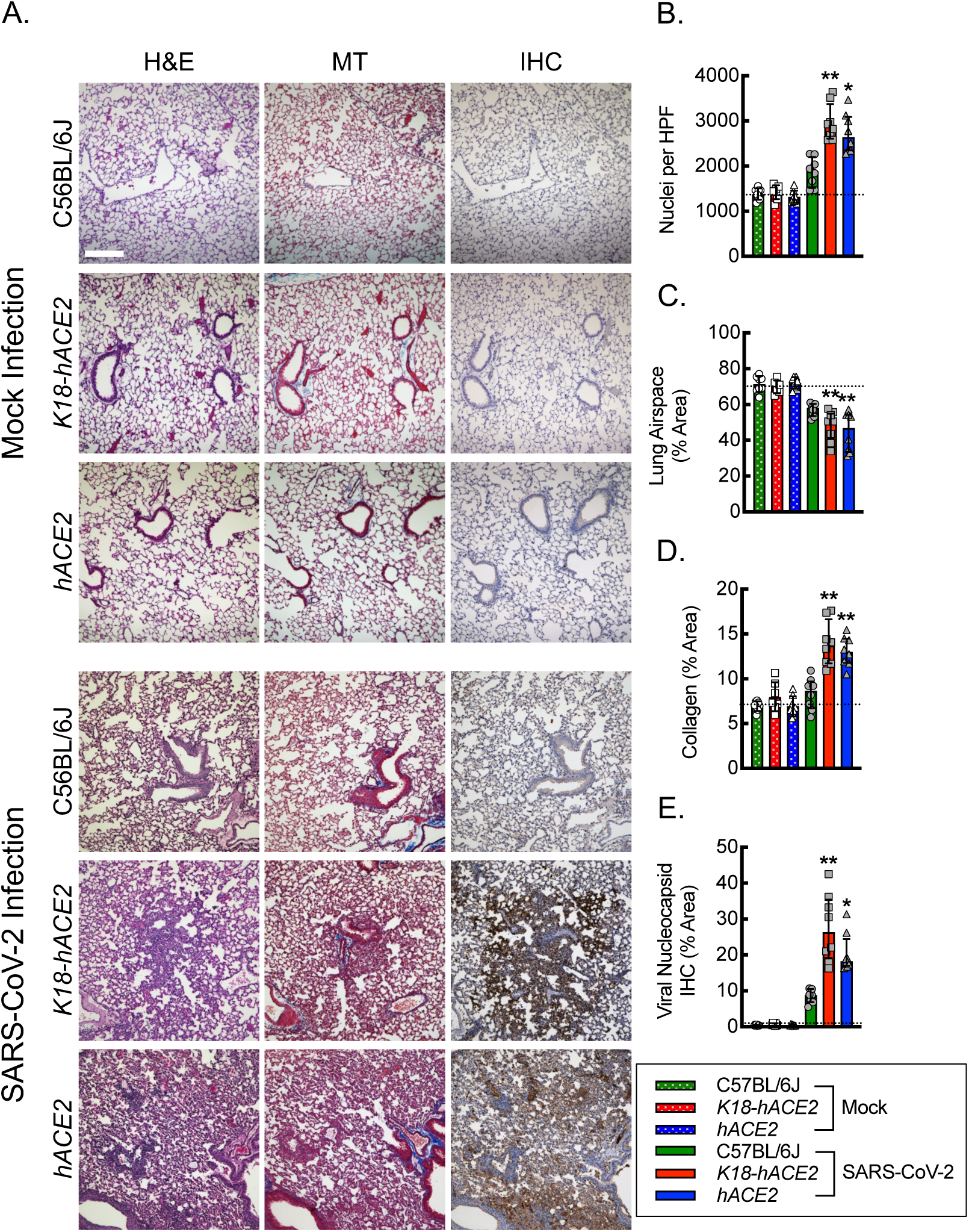
Lung tissue from *hACE2* mice infected with SARS-CoV-2 exhibits comparable measures of pneumonia relative to *K18-hACE2* mice. (A) Representative photomicrographs of lung histological sections on day 6 post intranasal infection with vehicle control (mock) or 1000 PFU/dose of SARS-CoV-2 infection stained with hematoxylin and eosin (H&E), Masson’s Trichrome (MT) and immunohistochemistry (IHC) for SARS-CoV-2 nucleocapsid protein. Bar, 200 microns. Quantitation of photomicrographs from histological sections of H&E to define number of nuclei (B) and relative lung airspace (C), of MT to define relative area of collagen (D), and of IHC for relative area of SARS-CoV-2 Viral Nucleocapsid protein (E) (*, P<0.05; **, P<0.001; compared to mock or C57BL/6J controls using ANOVA; n=8, 4 male and 4 female). Dotted line, average negative baseline on mock controls. Data, mean ± SD.

### Plasma from *hACE2* mice infected with SARS-CoV-2 exhibits a unique cytokine profile relative to infected C57BL/6J or *K18-hACE2* mice

To determine whether there was a unique inflammatory biomarker signature exhibited by *hACE2* mice relative to *K18-hACE2* or C57BL/6J mice, we carried out extensive chemokine, cytokine, and growth factor profiling on plasma samples collected 6 days post-viral exposure (**Fig. 5A**). Quantitation of the panel revealed five distinct patterns. Pattern 1 consisted of chemokines, cytokines, and growth factors, including CCL5, CCL7, IL-6, Leptin, TNF-*α,* and CSF-3 that were significantly increased in *K18-hACE2* mice relative to C57BL/6J and *hACE2* (**Fig. 5B-G**). Pattern 2 revealed that CCL2 and IL-10 were significantly elevated in both the *K18-hACE2* and *hACE2* mice relative to C57BL/6J (**Fig. 5H, I**). Pattern 3 consisted of IL-1*α*, IL-1β, IL-17*α*, and IL-22, and all were significantly increased in *hACE2* mice relative to C57BL/6J and trended higher in *hACE2* mice than in *K18-hACE2* mice (**Fig. 5J-M**). Pattern 4 revealed that VEGF-A was significantly decreased in *K18-hACE2* mice relative to C57BL/6J and *hACE2*, both of which survive infection (**Fig. 5N**). Finally, pattern 5 revealed that IL-33R was significantly decreased in both *K18-hACE2* and *hACE2* mice relative to C57BL/6J, which did not exhibit any evidence of lung pathology (**Fig. 5O**).

**Figure 5.**
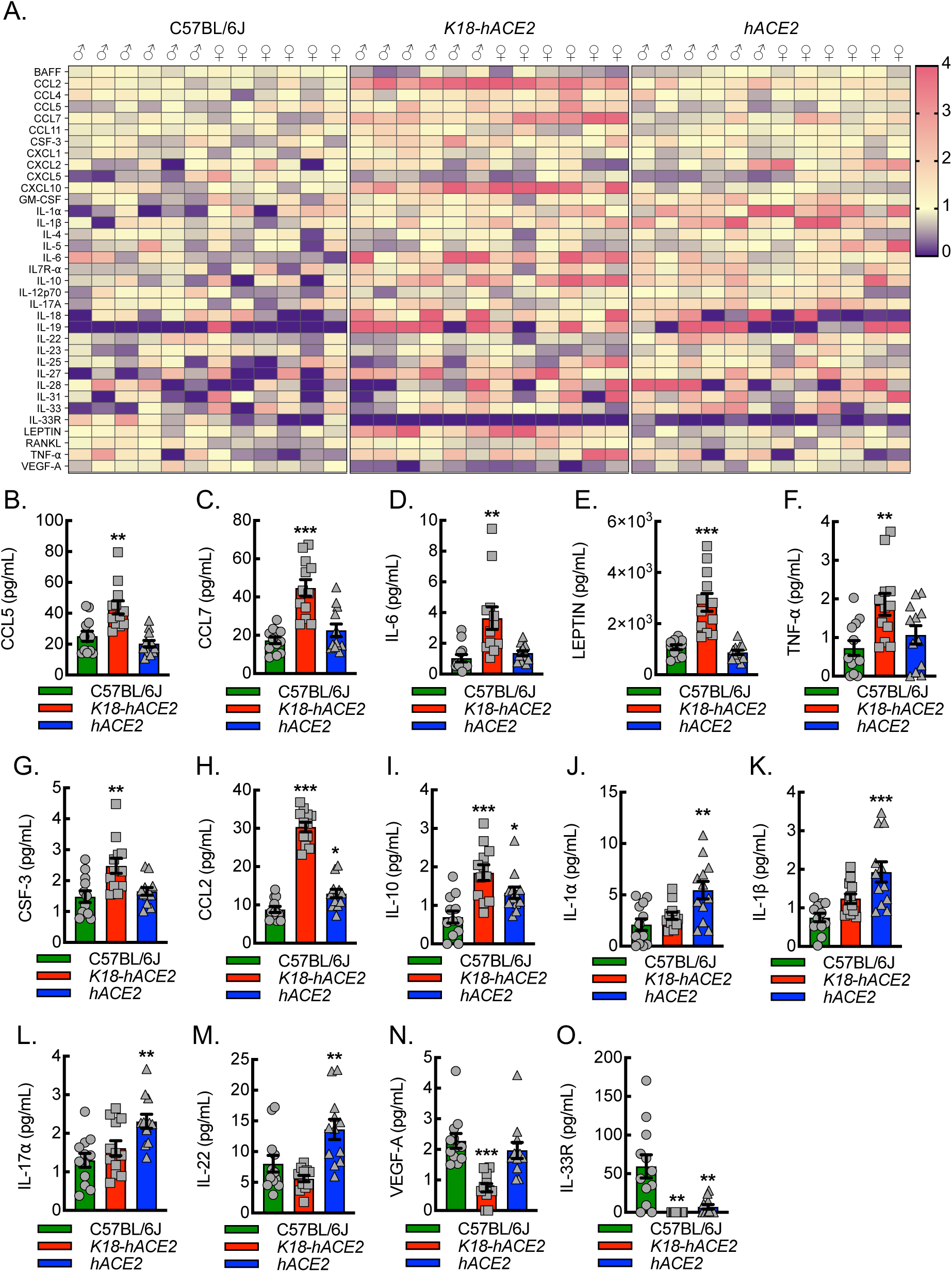
Plasma from *hACE2* mice infected with SARS-CoV-2 exhibits a unique cytokine profile relative to infected C57BL/6J or *K18-hACE2* mice. (A) Heat map depicting relative plasma inflammatory chemokine, cytokine, and growth factor expression patterns, as measured by Luminex® INTELLIFLEX, from C57BL/6J, *K18-hACE2*, or *hACE2* mice 6 days after intranasal infection with 1000 PFU/dose of SARS-CoV-2. Pattern 1 revealed elevations specific to *K18-hACE2* mice and included CCL5 (B), CCL7 (C), IL-6 (D), LEPTIN (E), TNF-α (F), and CSF-3 (G) (**, P<0.001; ***, P<0.0001 compared to all others using ANOVA; n=12, 6 male and 6 female). Pattern 2 revealed elevations in *K18-hACE2* and *hACE2* mice and included CCL2 (H) and IL-10 (I) (*, P<0.05; ***, P<0.0001 compared to C57BL/6J using ANOVA; n=12, 6 male and 6 female). Pattern 3 revealed elevations specific to hACE2 mice and included IL-1α (J), IL-1β (K), IL-17α (L), and IL-22 (M) (**, P<0.001; ***, P<0.0001 compared to all others using ANOVA; n=12, 6 male and 6 female). Pattern 4 revealed decreased expression specific to *K18-hACE2* mice and included VEGF-A (N) (***, P<0.0001 compared to all others using ANOVA; n=12, 6 male and 6 female). Pattern 5 revealed decreased expression in both the *K18-hACE2* and *hACE2* mice and included IL-33R (O) (**, P<0.001 compared to C57BL/6J using ANOVA; n=12, 6 male and 6 female).

### SARS-CoV-2 replicates more efficiently in macrophages of *hACE2* mice relative to *K18-hACE2* mice

Viral genomic replication is initiated by the synthesis of full-length negative-sense genomic copies, which function as templates for the generation of new positive-sense genomic RNA (63). Therefore, to assess whether lung macrophages were infected by the virus in the *hACE2* mice, we quantitated the colocalization of the negative strand of SARS-CoV-2 by fluorescence in situ hybridization (FISH) with macrophage cellular markers, CD68 (**Fig. 6**) and CD115 (**Fig. S4**). Our data demonstrated that *hACE2* mice recruit the largest number of CD68^+^ cells to the site of lung infection and that a significant number, 60-70%, co-stained positive for both CD68 and the SARS-CoV-2 negative strand. *K18-hACE2* mice revealed a 60% reduction in CD68^+^ cell recruitment to the site of infection relative to *hACE2* mice, and only about 25% of CD68^+^ cells co-stained positive for SARS-CoV-2 negative strand, though negative strand staining in the airways was quite evident in these mice. C57BL/6J mice demonstrated only a few CD68^+^ cells in the infected lung tissue with no evidence of SARS-CoV-2 negative strand positivity. The specificity of CD68 for macrophages has been called into question as some mesenchymal cells may also upregulate CD68 expression in response to inflammation (64, 65). To further support the specificity of detection in macrophages, we carried out immunofluorescence staining with an alternative and well-established myeloid marker, CD115 (colony-stimulating factor 1 receptor, Csf1r). Using CD115 in place of CD68 demonstrated comparable results regarding the localization of SARS-CoV-2 negative strand FISH, supporting the likelihood that the vast majority of CD68 positive cells reflect macrophages in this model. In summary, *hACE2* mice demonstrate evidence of viral replication in infiltrating macrophages relative to infected *K18-hACE2* and C57BL/6J mice, exhibiting a fundamental difference in the innate immune response to infection that may provide new insights into COVID-19.

**Figure 6.**
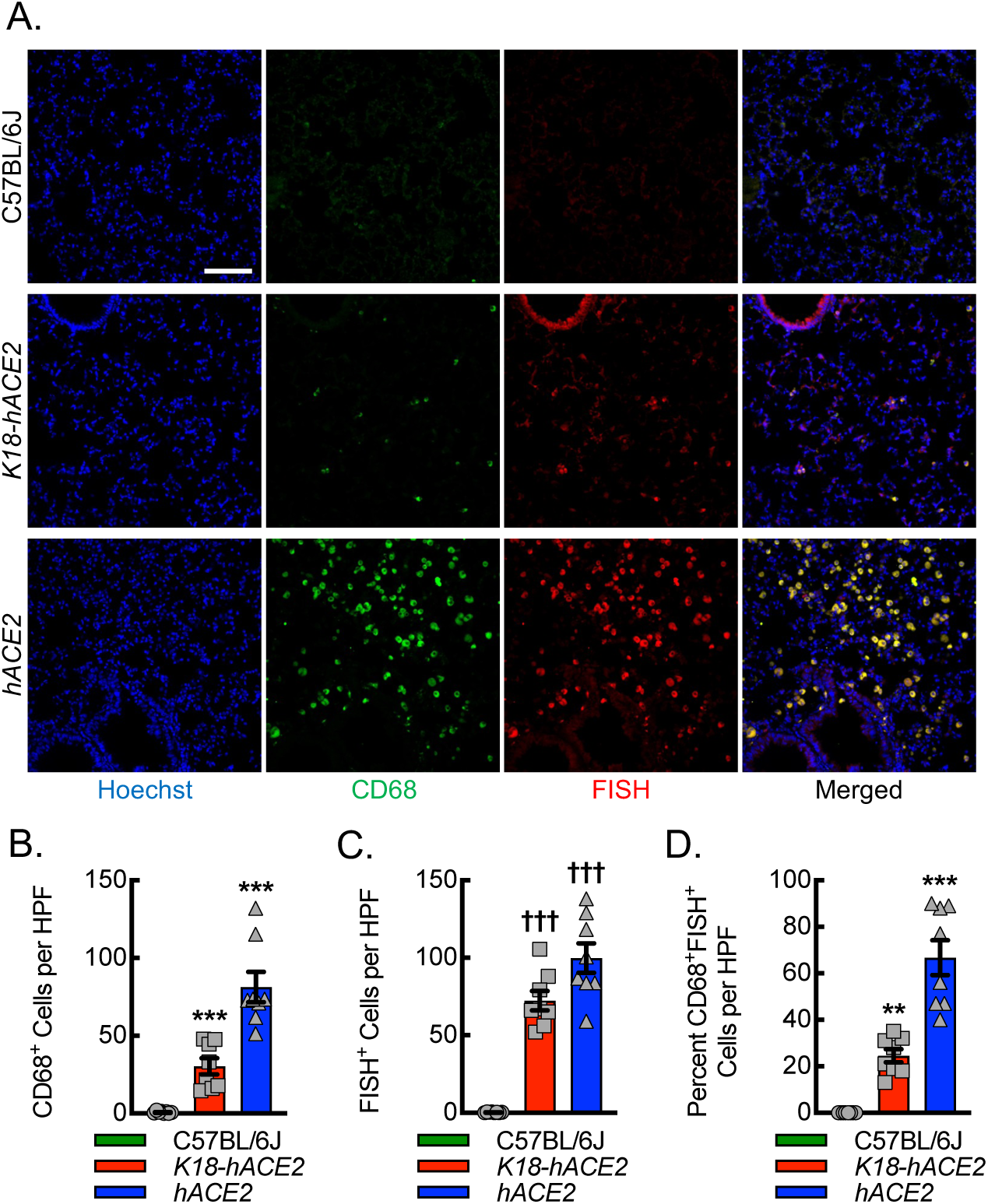
SARS-CoV-2 replicates more efficiently in macrophages of *hACE2* mice relative to *K18-hACE2* mice. (A) Representative immunofluorescent photomicrographs from lungs of C57BL/6J, *K18-hACE2*, and *hACE2* mice harvested on day 6 post intranasal infection with 1000 PFU/dose of SARS-CoV-2. Blue, Hoechst. Red, fluorescent in situ hybridization (FISH) for SARS-CoV-2 negative strand. Green, CD68. Bar, 100 microns. Quantitation of the number of CD68^+^ cells (B), number of FISH^+^ cells (C), and percent of CD68^+^ cells that are also FISH^+^ (D). (**, P≤0.009; ***, P<0.0001 compared to all others using ANOVA; ^†††^, P<0.0001 compared to C57BL/6J using ANOVA; n=8, 4 male and 4 female).

To confirm that CD68^+^ cells in the lungs of SARS-CoV-2 infected mice also demonstrated expression of hACE2 protein, we quantitated the number of CD68^+^ cells, the number of hACE2^+^ cells, and the percent of CD68^+^ cells that were also hACE2^+^ (**Fig. S5**). We found that about 88% of cells expressing CD68 also co-expressed hACE2 in the *hACE2* mice and that this was decreased by about 30% and 85% in the *K18-hACE2* and C57BL/6J mice, respectively. This confirmed that CD68^+^ cells in the *hACE2* mice infected by SARS-CoV-2 were recruited to the lung and expressed detectable hACE2 protein levels.

### Macrophage ACE2 expression is regulated by IL-1β-dependent NF-𝛋B transcription, and transcriptomic analysis of inflammatory macrophages infected with SARS-CoV-2 reveals altered expression of noncoding RNA and ribosomal machinery

ACE2 is considered the primary receptor for the SARS-CoV-2 virus (45, 50). To define potential mechanisms of macrophage viral uptake, we set out to determine ACE2 expression in primary bone marrow-derived macrophages (BMDMs) from *hACE2* mice along with the impact of inflammasome activity on ACE2 expression. We crossed *hACE2* mice to a strain containing the allelic combination of B6(FVB)-*Il1b^em1Mora^*/J (*IL-1β^fl/fl^*) and FVB-Tg(Csf1r-cre/Esr1*)1Jwp/J (*Csf1r^mericremer^*), allowing for inducible knockdown of IL-1β in macrophages (mIL-1β KO) (66, 67). BMDMs from *hACE2* control and mIL-1β KO *hACE2* mice were subjected to strong inflammasome stimulation by the combination of lipopolysaccharide (LPS) priming and cholesterol crystal exposure. Control mice consequently demonstrated increased expression of *IL-1β* mRNA as well as increased secretion of mature IL-1β protein (**Fig. 7A, B**). BMDMs from mIL-1β KO *hACE2* mice demonstrated a 75% decrease in both *IL-1β* mRNA expression and mature IL-1β protein secretion. Of interest, the *ACE2* mRNA expression pattern mirrored the *IL-1β* mRNA expression pattern, demonstrating the highest mRNA levels with the combination of LPS priming and cholesterol crystal exposure (**Fig. 7C**). The *IL-1β*-deletion led to a consequential knockdown in *ACE2* mRNA expression, supporting that macrophage ACE2 expression is dependent on IL-1β expression and autocrine signaling.

**Figure 7.**
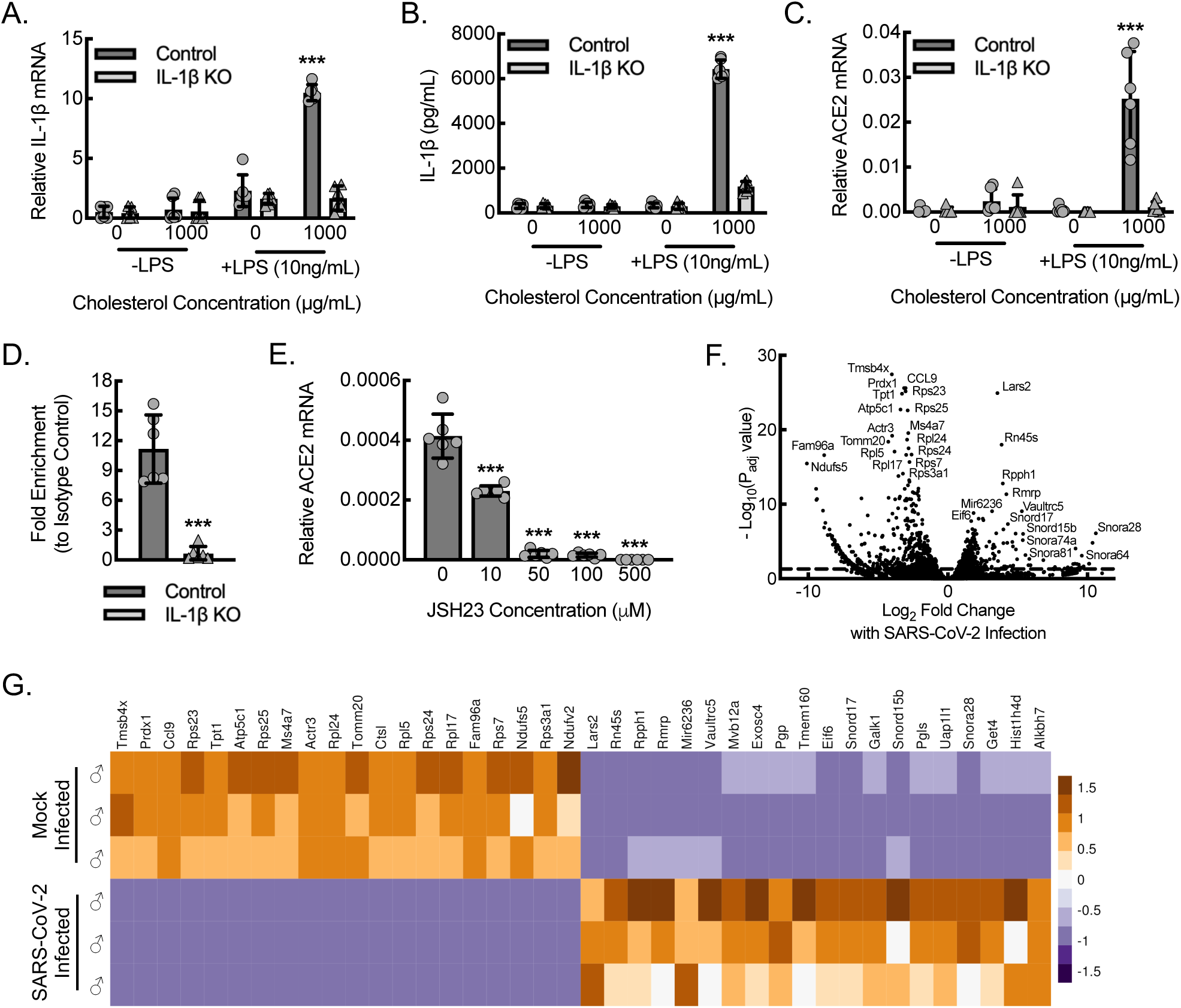
Macrophage ACE2 expression is regulated by IL-1β-dependent NF-𝛋B transcription and transcriptomic analysis of inflammatory macrophages infected with SARS-CoV-2 reveals altered expression of noncoding RNA and ribosomal machinery. (A) BMDMs from control or myeloid *IL-1β*-deleted (mIL-1β KO) *hACE2* mice were primed with or without LPS (10 ng/mL) and exposed to cholesterol crystals (0 or 1000 μg/mL) for 24 hours to induce inflammasome activation, followed by RT-qPCR for relative *IL-1β* mRNA expression (***, P<0.0001 compared to all others using ANOVA; n=6 mice total, 3 male and 3 female). (B) BMDMs treated as in (A), followed by ELISA on cell culture supernatants for secreted, mature IL-1β protein (***, P<0.0001 compared to all others using ANOVA; n=6 mice total, 3 male and 3 female). (C) BMDMs treated as in (A), followed by quantitative RT-qPCR for relative *hACE2* mRNA expression (***, P<0.0001 compared to all others by ANOVA; n=6 mice total, 3 male and 3 female). (D) Quantitative PCR of the *ACE2* promoter after ChIP, using antibody specific to p65 (RELA) relative to isotype control, of lysate from BMDMs stimulated with LPS+IFN-γ for 24 hours (***, P=0.0001 by *t*-test; n=6 mice total, 3 male and 3 female). (E) BMDMs from control or mIL-1β KO mice were treated with LPS+IFN-γ for 24 hours in the presence or absence of the NF-κB inhibitor, JSH23, at indicated concentrations, followed by RT-qPCR for relative *hACE2* mRNA expression in cell lysates (***, P≤0.0001 compared to control at 0 concentration by ANOVA; n=6 mice total, 3 male and 3 female). (F) Volcano plot demonstrating differential RNA-seq transcriptome data from BMDMs cultured from *hACE2* mice and stimulated with LPS+IFN-γ for 18 hours, followed by infection with SARS-CoV-2 for 24 hours (n=3 male mice). Dashed line, P_adj_=0.05. (G) Heatmap illustrating normalized counts for the top 20 differentially expressed genes derived from (F) with each column representing an individual mouse and each row representing mRNA from a single gene (n=3 male mice). Data, mean ± SD.

We previously demonstrated that nuclear activity of the p65 component of NF-κB in macrophages was dependent on IL-1β expression (66). We therefore took a candidate-based approach and performed a chromatin immunoprecipitation (ChIP) assay on BMDMs stimulated by the combination of LPS and interferon-gamma (LPS+IFN-γ), using antibodies specific to the p65 (RELA) component of NF-κB, to confirm IL-1β-dependent binding of this transcription factor to the ACE2 promoter region (**Fig. 7D**). Quantitative PCR with primers directed at the regions of the *ACE2* promoter associated with established p65 seed sites demonstrated that control BMDMs stimulated by LPS+IFN-γ achieved about 3-fold increased promoter binding using NF-κB-specific antibody relative to the respective isotype control antibody. Moreover, BMDMs from mIL-1β KO mice stimulated by LPS+IFN-γ demonstrated no increase over isotype control, supporting that NF-κB binds to the *ACE2* promoter in an IL-1β-dependent manner. To assess the functional effect of NF-κB binding to the *ACE2* promoter region, we treated LPS+IFN-γ-stimulated BMDMs with JSH23, an inhibitor of NF-κB. Inhibitor dose ranges were selected based on the established IC50s and our prior work (66, 68). Wild-type BMDMs stimulated by LPS+IFN-γ were treated with increasing concentrations of JSH23 and demonstrated dose-dependent reductions in *ACE2* mRNA expression, indicating that ACE2 expression is upregulated by IL-1β-dependent NF-κB activity (**Fig. 7E**).

Next, we assessed whether phenotypically inflammatory BMDMs (polarized by LPS+IFN-γ) from *hACE2* mice exhibited a response to SARS-CoV-2 infection in vitro, using a multiplicity of infection of 1.0 of SARS-CoV-2 followed by differential RNA-seq transcriptomic profiling (**Fig. 7F, G**). SARS-CoV-2 infected macrophages led to increased expression of members belonging to several noncoding RNA families: 1) small nucleolar RNAs (snoRNAs), Snord17, Snord15b, Snora28, Snora74, Snor81, and Snor64; 2) Vault RNA (vtRNA), Vaultrc5; 3) ribosomal RNA, Rn45s; 4) mitochondrial endoribonuclease RNA, Rmrp; and 5) microRNA, miR6336. SARS-CoV-2 infection of inflammatory macrophages also led to downregulation of several mRNAs encoding ribosomal protein components, including Rps23, Rps25, Rpl24, Rpl5, Rps24, Rpl16, and Rps3a1. Gene ontology analyses revealed these expression changes to be consistent with altered structural constituents of the host ribosome and consequently ribosomal biogenesis through both up- and downregulation of several components, impacting cytoplasmic translation and host mRNA binding (**Fig. S6**). Gene ontology also identified alterations in pathways potentially relevant to innate defense, including upregulation of autophagy, alterations in mitochondrial structure and function, including cellular respiration, and alterations in oxidation-reduction regulatory pathways.

## DISCUSSION

Here, we validated an experimental mouse model of COVID-19 in which *hACE2* replaces *mACE2* and is expressed under regulation of the native mouse promoter. While *hACE2* mice demonstrated susceptibility to SARS-CoV-2 infection with clear histological evidence of pneumonia, they exhibited a disease course that differs from the commonly used *K18-hACE2* mouse model in several ways that may help provide new mechanistic insights into the human disease pathobiology. One of the challenges of recapitulating aspects of human COVID-19 is its heterogeneity in clinical presentation with the vast majority of cases being minimally symptomatic and case fatality considered to be under 1% globally (1, 69). Experimental COVID-19 models have made use of either transgenic mice overexpressing human ACE2 like the *K18-hACE2* mice, mouse-adapted strains of the SARS-CoV-2 virus, or more recently, mice with humanized immune systems that require adenoviral-induced overexpression of human ACE2 to induce infection susceptibility (58, 70–72). The *K18-hACE2* mice are well-established and known to exhibit efficient *hACE2* transgene overexpression in the airway epithelial cells but not alveolar epithelia and in the epithelia from other organs like the central nervous system, gastrointestinal tract, and kidneys (58, 61, 73, 74). This differs somewhat from native mouse or human expression of ACE2, which is quite dynamic and ubiquitous, including expression in the cardiovascular and respiratory systems, central nervous system, gastrointestinal system, renal system, and adipose tissue, where it functions to regulate the renin-angiotensin system and to facilitate amino acid transport (75). Following intranasal challenge, *K18-hACE2* mice develop severe pneumonia characterized by suppurative rhinitis, alveolar necrosis, edema, hemorrhage, fibrin deposition, alveolar and interstitial inflammatory infiltrates, and vasculitis with thrombosis (60–62). The *K18-hACE2* model exhibited elevations in both local and systemic chemokine and cytokine expression in a manner analogous to the cytokine profiles associated with the development of ARDS and extrapulmonary multi-organ failure in human disease. Infections in *K18-hACE2* mice can be highly lethal (>90% mortality by 6-7 days of infection), making them an attractive model for acute fulminating disease but limiting the ability to study mechanistic determinants of varying severity as well as long-term outcomes. The overall global case fatality for human COVID-19 has decreased over time with the evolution of the virus to the extent that mortality is far lower than what is represented in the *K18-hACE2* model (76). The emergence of an endemic SARS-CoV-2 cycle has also raised concern for an increasing prevalence of patients who experience PASC with longer-term functional limitations (10). Comparable to *K18-hACE2* mice, histopathology of lung tissue from *hACE2* mice 6 days post-infection supported evidence of pulmonary consolidation that exhibited cellular infiltrate, edema, collagen deposition, and alveolar hemorrhage. Relative to the *K18-hACE2* model, the *hACE2* mice with experimental COVID-19 exhibited a presentation somewhat more analogous to most human cases, as evidenced by maintenance of stable weight and survival, making them ideally suited for studying determinants of severity and long-term outcomes.

Several factors appear to be critical for successful SARS-CoV-2 clearance and resolution of clinical infection, including virulence properties of the strain, dose and route of exposure, environment, and genetic and immune properties of the host (37). Dynamic cellular expression of ACE2, given its role as a major receptor for the virus, may influence or be influenced by such factors (46–48). As noted, human ACE2 and mouse ACE2 proteins are homologous, but mouse ACE2 lacks efficient SARS-CoV-2 binding due to key amino acid substitutions in the SARS-CoV-2 spike protein binding domain (44, 45, 49). Human and mouse *ACE2* promoter sequences contain evolutionarily conserved regions upstream of the *ACE2* transcriptional start site (54, 55). Thus, a model of native promoter regulation in which mice survive may provide mechanistic insights into aspects of host and immune response that other models, by their design, are unable to provide.

There is tremendous interest in using cytokine expression patterning as a predictive biomarker for the severity of disease. Yet, our understanding of the kinetics of systemic cytokine expression associated with the widely varied clinical presentations of COVID-19 remains incomplete (25). Cytokine profiling of SARS-CoV-2-infected *hACE2* mice revealed a unique signature relative to C57BL/6J and *K18-hACE2* mice, and it is interesting to speculate whether the patterns may be providing clues to the severity of infection or likelihood of viral clearance and survival. *K18-hACE2* mice, which exhibited high mortality at the doses of SARS-CoV-2 used in this study, demonstrated significant elevations in plasma chemokines (CCL5 and CCL7), proinflammatory cytokines (IL-6, Leptin, and TNF-α), and a growth factor (CSF-3), relative to both C57BL/6J and *hACE2* mice. Both *K18-hACE2* mice and *hACE2* mice exhibited elevations in CCL2 and IL-10 relative to C57BL/6J, yet plasma levels in *K18-hACE2* mice were higher. Elevated levels of CCL2 and CCL7, two chemokines potent at the recruitment of CCR2^+^ monocytes, have also been found in bronchoalveolar lavage fluid from patients with severe COVID-19 (77). In hospitalized patients, individuals requiring intensive care unit (ICU) level of care demonstrated higher systemic levels of the inflammatory cytokines and chemokines: IL-2, IL-7, IL-10, G-CSF, IP-10, CCL2 (MCP-1), CCL3 (MIP-1α), and TNF-α (1). *hACE2* mice, which exhibited survival, demonstrated significant elevations in plasma IL-1α, IL-1β, IL-17α, and IL-22 levels. In separate work, we have demonstrated that IL-1β is critical for expression of VEGF-A to set the stage for vascular regeneration and healing (66). Both IL-1β and VEGF-A have been described as broadly elevated in hospitalized COVID-19 patients, irrespective of ICU level of care (1). IL-17α is known to serve in both proinflammatory and anti-inflammatory roles by, in part, increasing the local production of chemokines to promote recruitment of monocytes/macrophages as well as other immune cells (78, 79). IL-22 can target epithelial and stromal cells where it can promote proliferation and play a role in tissue regeneration (80). Both C57BL/6J and *hACE2* mice exhibited elevation in the growth factor VEGF-A relative to the *K18-hACE2* mice, suggesting an association of VEGF-A with survival in these models. Finally, C57BL/6J mice, which did not exhibit overt clinical pathology despite detectable viral load, demonstrated increased levels of IL-33R relative to either *hACE2* or *K18-hACE2* mice. It is interesting to note that IL-33R is expressed by several cell types in the lung and plays a role in myeloid differentiation and epithelial repair (81).

Another key difference between SARS-CoV-2-infected *hACE2* and *K18-hACE2* mice was the evidence of increased SARS-CoV-2 replication in lung macrophages of the *hACE2* mice. The mechanism of macrophage cell infection appeared to be linked to dynamic ACE2 expression and depended upon IL-1β-driven NF-κB transcription of *hACE2*. Macrophages are established innate sensors of danger signals from microbial pathogens via pattern recognition receptors (PRRs), releasing inflammatory molecules that eliminate pathogens, initiate inflammation and recruitment of additional effector cells, and promote tissue repair (34). It is currently unclear whether the SARS-CoV-2 replication in macrophages represents a component of this innate response ultimately promoting clearance of the virus, tissue repair, and healing, but it is interesting to note the correlation between increased infection of macrophages and the overall survival in the *hACE2* mice relative to the *K18-hACE2* model. Dysregulated macrophage responses and macrophage activation syndrome identified early in the COVID-19 pandemic were associated with hyperinflammation and were thought to pose a substantial danger for tissue along with increased susceptibility to secondary bacterial superinfection (22, 34, 82–85). Several viruses use monocytes and macrophages as vessels for replication, dissemination, or long-term persistence within tissues, inducing the expression of proinflammatory signaling and antiviral molecules (86–91). So, it remains possible that infection of macrophages or infection of macrophages that are dysfunctional from other comorbidities (i.e. age, cardiovascular disease, diabetes mellitus) may contribute to more severe infection and worsening clinical presentation. Direct infection of macrophages with SARS-CoV-1 led to the induction of several chemokines that amplified inflammation but with an impaired type I interferon (IFN) response, potentially worsening tissue damage with consequent impaired viral clearance (92). Comparable concerns regarding interferon response in COVID-19 have also been raised (19, 20). More work is required in the study of macrophage infection responses during experimental COVID-19. Still, it appears that *hACE2* mice may provide a model to facilitate complementary mechanistic studies in the context of comorbidities like aging, experimental atherosclerosis, or diabetes.

Competition between viral usurpation of RNA processing machinery and the cellular innate antiviral defenses is exhibited by cells during infection with many RNA viruses, including influenza, human immunodeficiency virus type-1 (HIV-1), dengue, rabies, and SAR-CoV-2 (93–97). Compared to mock-infection, SARS-CoV-2 infection of inflammatory macrophages in vitro led to a significant alteration in the RNA expression profile. The upregulation of several noncoding RNAs, including Rn45s, multiple snoRNAs, and Vaultrc5, is suggestive of an attempt to co-opt ribosomal RNA machinery to help with viral replication. Rn45s serves as a precursor for the 18S, 5.8S, and 28S rRNA subunits and is required for ribosomal biogenesis (98). SnoRNAs are highly expressed non-coding RNAs that play an important role in binding and processing of rRNA, and knock-down studies have demonstrated that RNA viruses can usurp snoRNAs for replication (94, 99). Vaultrc5 is a member of the highly conserved Vault small non-coding RNA family, known to be increased in expression in response to RNA viruses to support replication (97, 100). Moreover, downregulation of several mRNAs encoding ribosomal protein components is indicative of an attempt to alter the structural constituents of the host ribosome, which can lead to inhibition of host mRNA binding as well as impaired cytoplasmic translation of host proteins (96). Finally, pathways important for the innate immune response to the virus-like autophagy, mitochondrial cellular respiration, and oxidation-reduction were identified by gene ontology as upregulated, indicating activation of macrophage antiviral defenses (101–104).

There are several limitations to this study. In addition to increased levels of acute phase reactants and pro-inflammatory cytokines, severe COVID-19 presentations have also been associated with peripheral immune activity, neutrophilia and emergence of immature and low-density neutrophils, increased neutrophil-to-lymphocyte ratio and lymphopenia, myeloid inflammation, and reduced expression of the human leukocyte antigen DR isotype (HLA-DR) by circulating monocytes (18, 25, 31, 105–110). To validate *hACE2* mice as a potential experimental model of COVID-19, we chose to focus on a few key assessments in comparison with C57BL/6J and *K18-hACE2* mice, including survival, weight loss, histopathology, cytokine profiling, and evidence of infection/viral replication in lung macrophages. We did not determine whether macrophage infection plays a role in pathology or facilitates viral clearance. Future studies involving the impact of macrophages on disease course as well as broader aspects of the immune response, will provide additional insight into the clinical relevance of the pathways identified by this model.

Additionally, assessment of *hACE2* mice under conditions of altered innate immune response like aging, vascular disease, and diabetes may permit investigation of mechanisms determining differences in severity of pathology and further elucidate the role of macrophage infection. It is known that patients requiring ICU-level care were older and had a higher prevalence of hypertension, cardiovascular disease, and diabetes (111). In the assessment of variations in clinical presentation, it will be important to include additional physiological parameters like heart rate, temperature, blood pressure, and pulse oximetry to obtain a more complete clinical picture. While *hACE2* mice survived infection at a SARS-CoV-2 dose that was lethal to most *K18-hACE2* mice, it remains possible that *hACE2* mice may exhibit mortality with higher doses, and thus it will be important to carry out additional dose-response curves in the future. If a heterogeneous syndrome like PACS is to be studied in the *hACE2* mice model, the appropriate prioritization of metrics that can assess the complement of long-term impairments will be required. Finally, it will be important to validate the translational relevance of our mechanistic findings utilizing human specimens.

In conclusion, we have developed a model of COVID-19 by humanizing ACE2 expression under regulation of the native mouse promoter to determine whether macrophages may be infected by SARS-CoV-2. Mice demonstrate histopathologic evidence of pneumonia upon intranasal exposure to SARS-CoV-2, and they survive and maintain normal weight relative to the previously established *K18-hACE2* model, which exhibits severe weight loss and high mortality. Plasma chemokine, cytokine, and growth factor profiling of *hACE2* mice during infection reveals a unique pattern associated with aspects of an inflammation, regeneration, and healing signature that supports survival. Finally, evidence of infection and viral replication within macrophages was more prevalent in the *hACE2* mice relative to *K18-hACE2* mice. Thus, *hACE2* mice may serve to model appropriate and inappropriate innate immune responses in the context of COVID-19. Native promoter regulation of ACE2 expression in macrophages appears dynamic and based upon IL-1β-driven NF-κB transcription of ACE2, and this mechanism may be potentially targeted to influence the macrophage response to infection. The *hACE2* mice presented here may provide additional mechanistic insights into transcriptional regulatory elements that promote variation in ACE2 expression and consequently variation in COVID-19 presentation and severity. Adequate disease modeling may help to define underlying innate immune cell mechanisms that determine the variable severity of disease presentation and thereby yield druggable targets that ameliorate risk associated with the more severe forms of this disease.

## ACKNOWLEDGMENTS

Research reported in this publication was supported by the Harold S. Geneen Charitable Trust Awards Program for Coronary Heart Disease Research (A.R.M.), and by Research Project Grants NIH NHLBI R01HL139795 (A.R.M.), R01HL163005 (A.R.M.), R01HL148727 (G.C.). This work was supported by IDeA NIH NIGMS P20GM103652 (A.R.M., C.S.M., and G.C.) and P30GM149398 (G.C.). This work was supported by VA VHA CSR&D 1I01CX002231 (A.R.M.), VA VHA CSR&D I01CX001892 (G.C.). This work was supported by a Biocontainment Research Support Services Core NIH UC7AI180306 (R.O.). This work was supported by AHA 23CDA1056587 (C.S.M.) and RI Foundation #16415_139169 (C.S.M.). This work was also supported by NIH T32HL134625 Brown Respiratory Research Training Program (R.C.), and R25HL088992 (C.P.) and by NIH NIGMS 8P30 GM103410 (The Mouse Transgenic and Gene Targeting Facility at Brown University). The views expressed in this article are those of the authors and do not necessarily reflect the position or policy of the Department of Veterans Affairs or the U.S. government. The funders had no role in study design, data collection and analysis, decision to publish, or preparation of the manuscript.

## AUTHOR CONTRIBUTIONS

A.R.M., G.C., and R.M.O. conceived the study. C.L., R.K., C.S.M., S.S., J.P., E.A., C.A.B., A.F., C.P., O.C., J.A.M., S.U., J.J.W., J.L.K., X. Z., G.C., R.M.O., and A.R.M. performed the in vitro and animal experiments. C.L., R.K., C.S.M., S.S., J.P., E.A., C.A.B., A.F., C.P., O.C., J.A.M., S.U., J.J.W., J.L.K., X. Z., G.C., R.M.O., and A.R.M. analyzed the data. C.L., R.C., C.S.M., J.P., C.A.B., J.L.K., X.Z., G.C., R.M.O., and A.R.M. wrote the manuscript.

## INCLUSION AND DIVERSITY

We worked to ensure sex balance in the selection of non-human subjects. One or more of the authors of this paper self-identifies as an underrepresented ethnic minority in science. One or more of the authors of this paper self-identifies as a member of the LGBTQ+ community. One or more of the authors of this paper self-identifies as living with a disability. One or more of the authors of this paper received support from a program designed to increase minority representation in science. While citing references scientifically relevant for this work, we also actively worked to promote gender balance in our reference list.

## METHODS

### RESOURCE AVAILABILITY

#### Lead Contact

Further information and requests for resources and reagents should be directed to and will be fulfilled by the lead contact, Alan R. Morrison.

#### Materials Availability

The *IL-1β^fl/fl^* mouse strain (JAX, 037836) (66) has been donated to The Jackson Laboratory consistent with NIH and VA resource sharing plans. *hACE2* mouse lines generated in this study will be made available upon request to the lead contact through use of a material transfer agreement (MTA) with Providence VA Medical Center. The MTA will be necessary to guide 1) general terms on how the mice will be used; 2) liabilities; 3) husbandry and shipping costs; 4) prevention of inappropriate distribution; 5) appropriate acknowledgement of the source; and 5) insurances that all planned animal subjects use is compliant with NIH and VA regulations.

#### Data and code availability

The lead contact will share data reported in this paper upon request. This paper does not report original code. Any additional information required to reanalyze the data reported in this paper is available from the lead contact upon request.

### EXPERIMENTAL MODEL AND SUBJECT DETAILS

#### Animals

C57BL/6J (JAX, 000664), *B6.Cg-Tg(K18-ACE2)2Prlmn/J* (*K18-ACE2;* JAX, 034860)(58), and FVB-Tg(Csf1r-cre/Esr1*)1Jwp/J (*Csf1r^mericremer^*; JAX, 019098) (67) mice were obtained commercially from The Jackson Laboratory. *IL-1β^fl/fl^* mice (66) were generated and validated previously by our group. The *hACE2* mice were generated by CRISPR/Cas9 technology by Brown University’s Mouse Transgenic and Gene Targeting Facility. Briefly, two recognized guide sites in exon 2 of the native *mACE2* locus that were conserved between mice and humans were identified. After the CRISPR/Cas9 system introduced a double-strand break in two targeted sites of the native *mACE2* allele, the homologous donor vector, composed of *hACE2* cDNA (Ace2-202, ENSMUST00000112271.9) with an SV40 polyA tail and flanked by homology arms, recombined, resulting in the insertion of the human *ACE2* cDNA into the exon 2 locus. The *hACE2* mice were bred in-house at the Providence VA Medical Center and shipped to the University of Missouri for acclimation before the challenge. For SARS-CoV-2 infections, mice 12-14 weeks of age were used. For BMDM isolations, animals 10-12 weeks of age were used. Male and female animals were used in equal numbers, and sex was analyzed as a biological variable by 2way ANOVA. All experiments were performed in accordance with the Providence VA Medical Center and the University of Missouri guidelines, and the Institutional Animal Care and Use Committee approved all housing protocols and experimental animal procedures. All animal procedures complied with the Office of Laboratory Animal Welfare and the National Institutes of Health Guide for Care and Use of Laboratory Animals. All mice used in these studies were reared at Providence VA Medical Center or the University of Missouri in AALAAC-approved animal housing facilities.

#### Cells and Virus

Vero E6 (ATCC, CRL-1586) cells were maintained at 37°C in high glucose Dulbecco’s modified Eagle medium (DMEM; Gibco, 11965092) supplemented with 10% serum plus II (Sigma-Aldrich, 14009C) and 1X GlutaMAX (Gibco, 35050061). The SARS-CoV-2 isolate USA_WA1/2020 was obtained from BEI resources (NR-52281). SARS-CoV-2 was propagated in Vero E6 cells according to BEI recommendations. Supernatant from infected Vero E6 cells was clarified by centrifugation, aliquoted for single-use, and stored at −80°C. All work with SARS-CoV-2 was performed in a biosafety level 3 laboratory by trained personnel equipped with powered air-purifying respirators. All experimental samples were processed by an approved protocol to inactivate any SARS-CoV-2 before removal from the biosafety level 3 laboratory or were analyzed at biosafety level 3.

#### SARS-CoV-2 Animal Infections

For challenge experiments, equal numbers of male and female age-matched mice (10-12 weeks) were transferred to the Laboratory for Infectious Disease Research (secure BSL-3 facility). Mice were anesthetized with 2-3% inhaled isoflurane and challenged by intranasal instillation of two 15μL droplets (one droplet per naris, 30μL total, equaling 1000 PFU/dose) of viral stocks diluted in sterile PBS or dilutions as indicated by figure legends. All infected mice were monitored by daily assignment of health scores, which involved assessments of appearance and activity, and weighed during the acute phase of the disease (14 days post-infection). For studies over 14 days, animals were monitored daily, weighed, and assessed weekly following the acute phase. Animals that survived to the end of the observation period or were identified as moribund (defined by pronounced neurologic signs, inactivity, severe weakness, or severe weight loss) were euthanized by CO_2_ asphyxiation followed by bilateral pneumothorax or cervical dislocation, methods approved by the American Veterinary Medical Association Guidelines on Euthanasia.

At the indicated day(s) post-infection, mice were euthanized, and blood and tissues were collected. Blood was collected in EDTA microtainers and further processed to separate plasma and buffy coats using SepMate tubes (STEMCELL Technologies, 85415) and Ficoll-Paque PREMIUM 1.084 (Cytiva, 17544602) according to the manufacturers’ instructions and immediately stored at −80°C until analysis. For nucleic acid extraction, tissues and buffy coat were placed in TRIzol™ (Fisher Scientific, 15596026) for SARS-CoV-2 inactivation. For histological analysis, tissues were placed in 10% neutral buffered formalin for at least 7 days for SARS-CoV-2 inactivation.

#### Quantitative RT-PCR

Total RNA from lung tissue (100-150 mg) was extracted using the TRIzol™. RNA was then column purified according to the manufacturer’s protocol (QIAGEN, RNeasy Kit 74106). RNA concentration was measured by Nanodrop One (Thermo Fisher). First-strand complementary DNA was synthesized with iScript cDNA synthesis Kit (Bio-Rad, 1708891). Each reaction used equal amounts of RNA (200ng/µl) as templates. Quantitative RT-PCR was performed with SSoAdvanved UnivSYBR Supermix (Bio-Rad, 1725274) on the StepOne Plus PCR machine (AB Applied Biosystems). All measurements were made in triplicate and then averaged, using primers directed at HPRT or GAPDH as the housekeeping genes. Primers: HPRT, 5’-GACCGGTCCCGTCATGCCGA-3’ (sense) and 5’-TGGCCTCCCATCTCCTCCATGACA-3’ (antisense); GAPDH, 5’-GTGTGAACGGATTTGGCCG-3’ (sense) and 5’-GTGATGGGCTTCCCGTTGAT-3’ (antisense); IL-1β, 5’-AAAGATGAAGGGCTGCTTCC-3’ (sense) and 5’-GTCCACGGGAAAGACACAGG-3’ (antisense); hACE2, 5’-TGAAACATACTGTGACCCCGC-3’ (sense) and 5’-TCCTCCTCCAACAGTTTCTGTCC-3’ (antisense); mACE2, 5’-GGATACCTACCCTTCCTACATCAGC-3’ (sense) and 5’-CTACCCCACATATCACCAAGCA-3’ (antisense); mACE1, 5’-CAGAATCTACTCCACTGGCAAGGT-3’ (sense) and 5’-TCGTGAGGAAGCCAGGATGT-3’ (antisense). Relative viral RNA was detected using primers specific for ORF1, S, and N (Thermo Fisher, A50137; Vi07921935_s1, Vi07918636_s1, Vi07918637_s1, respectively).

#### Immunoblot Analysis

Recombinant mouse (R&D Systems, 3437-ZN-010) or human (Abcam, ab151852) ACE2 protein (25 ng per lane) were collected with Laemmli sample buffer (Bio-Rad, 1610737) containing 2-mercaptoethanol (0.35 % v/v) and then boiled at 95°C for 5 minutes. Soluble proteins were separated by Tris/Glycine/SDS–PAGE (Bio-Rad, 1610732) and blotted into PVDF membranes (Bio-Rad, 1704157). Membranes were blocked in fluorescent blocking buffer (Rockland Immunochemicals, MB070) and probed at 4°C overnight with specific antibodies directed against human ACE2 (0.4 μg/mL; R&D Systems, MAB10823) or Mouse ACE2 (0.25 μg/mL; R&D Systems, AF3437) or a species nonspecific ACE2 antibody (0.4 μg/mL; R&D Systems, AF933). Secondary antibodies included Donkey anti-Rabbit IgG (H+L) Highly Cross-Adsorbed Secondary Antibody, Alexa Fluor™ 680, Invitrogen™ (0.5 μg/mL; ThermoFisher, A10043), Rabbit anti-Goat IgG (H+L) Cross-Adsorbed Secondary Antibody, Alexa Fluor™ 680 (0.5 μg/mL; ThermoFisher, A21088), IRDye® 800CW Donkey anti-Goat IgG Secondary Antibody (0.25 μg/mL LI-COR, 92632214). Immunoblots underwent imaging with the Odyssey CLX Imaging System (LI-COR). Relative species-specific antibody detection of ACE expression was quantitated by densitometry and normalized to non-specific antibody, using the Image Studio Lite (LI-COR, v1.0.18) or Image J software (112).

#### Cytokine analysis

Cytokines were assessed using a multiplex cytokine assay (ThermoFisher, EPX480-20834-901) on a Luminex Intelliflex instrument according to the manufacturer’s instructions with the following deviation: the cytokine assay was performed at biosafety level 3 up through the final incubation step. Paraformaldehyde (final concentration 4%) was then added to all wells (including standard wells) for the inactivation of SARS-CoV-2. The final wash, resuspension, and machine analysis steps were completed in a biosafety level 2 support lab.

#### Bone marrow-derived macrophage (BMDM) isolation and polarization

Using an established protocol, BMDMs were obtained by *in vitro* differentiation of primary femur and tibia bone marrow cells (113, 114). In brief, femurs and tibias from the indicated mouse lines mice were dissected, cleaned, disinfected in 70% ethanol, and washed with fully supplemented RPMI 1640 medium (Fisher Scientific, 11875085). Red blood cells were removed by ammonium-chloride potassium lysis buffer and subsequent centrifugation. The remaining BM cells were then cultured at a density of 3.5 × 10^6^ cells/10cm Petri dish in fully supplemented RPMI with 30% (vol/vol) L929 cell-conditioned medium. Cells were matured to phenotypic macrophages over 6-7 days as confirmed by >98% expression of F4/80 and CD68. Non-adherent cells were washed away with PBS, while adherent cells were recovered by gentle pipetting in PBS with 1mM EDTA. Polarization of BMDMs was carried out by incubation with LPS (100ng/mL; Sigma-Aldrich, L2630) and IFN-γ (50ng/mL; BioLegend, 575304) or with IL-4 (10ng/mL; BioLegend, 574304) and IL-13 (10ng/mL; BioLegend, 575904) up to 24 hours. In some experiments, BMDMs were treated with the NF-κB inhibitor, JSH23 (Abcam, ab144824).

#### Cholesterol crystal inflammasome assay

Cholesterol crystals were generated as previously described (115). Cholesterol (Sigma-Aldrich, C3045) was dissolved in 1-propanol at 2 mg/mL. Crystals were precipitated overnight at room temperature by dilution to 40% 1-propanol with water. Crystals were pelleted by centrifugation and dried at 70°C. Crystal pellets were then re-suspended in 0.1% FBS in PBS at a concentration of 50mg/mL. The average crystal size of 1-2μm was confirmed by microscopy. BMDMs (+/- priming with low dose (10ng/mL) LPS for 2 hours) were then either exposed to 1000μg/mL cholesterol crystals or vehicle control for 24 hours. Culture supernatants were collected, and IL-1β was quantified by ELISA.

#### Mouse IL-1β ELISA

Conditioned media from either polarized BMDMs or inflammasome-stimulated (cholesterol crystal assay) were used to evaluate mature IL-1β protein concentrations by ELISA (BioLegend, 432604), according to the manufacturer’s instructions. Each animal measurement reflects the average of experimental triplicates.

#### Chromatin Immunoprecipitation (ChIP) Assay

ChIP assay (Millipore, 17-295) was performed according to the manufacturer’s protocol. 1.5 x 10^6^ BMDMs were polarized by LPS and IFN-γ, followed by histone-DNA crosslinking with 37% formaldehyde. Cell debris and DNA were scraped and collected in PBS supplemented with protease inhibitors (1mM phenylmethylsulfonyl fluoride (ApexBio, A2587), 1μg/mL aprotinin (ApexBio, A2574), and 1μg/mL pepstatin A (ApexBio, A2571). The suspension was pelleted by centrifugation and then brought up in the SDS Lysis buffer supplied by the kit, supplemented with the protease inhibitors. The material was then sonicated with a Vibra-Cell™ sonicator (SONICS & Materials, VCX130) for 3 rounds of 15-second pulses and 5-second rest at 40% amplitude at room temperature. The sonicated cell supernatant was combined with ChIP Dilution Buffer (Millipore, 20-153) and Protein A/Salmon Sperm DNA beads (Millipore, 16-157C) and then incubated overnight at 4°C with antibodies directed at RelA (p65) protein (2.1μg/mL; Cell Signaling Technology, 8242) and its respective isotype control antibody (2.1 μg/mL; R&D Systems, AB-105-C). Additional Protein A Agarose/Salmon Sperm DNA beads were added to collect the DNA/histone complex, followed by centrifugation and further washing in a series of low- and high-salt buffers. Reversal of histone-DNA crosslinking was carried out by the addition of NaCl (5 M) for 4 hours at 65°C. DNA was obtained by phenol/chloroform extraction and ethanol precipitation. Site-specific primers for the ACE2 promoter region were used for quantitative PCR, and the results were depicted as cycle threshold fold enrichment relative to isotype control. ACE2 promoter primers: 5’-ATGGACAACTTCCTGACCGC-3’ (sense) and 5’-CTCAGGCTCATGATCTCGCC-3’ (antisense).

#### RNAseq analysis

In the secure BSL-3 facility, 3.0 x 10^6^ BMDMs were polarized by LPS and IFN-γ for 18 hours, followed by infection with SARS-CoV-2 dosed at an MOI of 0.1 for 24 hours. For RNA extraction, cells were collected and placed in TRIzol™ (Fisher Scientific, 15596026) for SARS-CoV-2 inactivation. RNA was extracted from TRIzol™ and then column purified (QIAGEN, RNeasy Kit 74106). Briefly, 2 μg of total RNA was processed at Active Motif (Carlsbad, California, USA) in Illumina’s TruSeq Stranded mRNA Library kit. Libraries were sequenced on Illumina NextSeq 500 as paired-end 42-nt reads. Sequence reads are analyzed with the STAR alignment—DESeq2 software pipeline(116–119). Briefly, reads generated by Illumina sequencing are mapped to the genome using the STAR algorithm with default settings. Alignment information for each read is stored in the BAM format. The nfcore/rnaseq pipeline is applied to perform the alignment. The number of fragments overlapping predefined genomic features of interest are counted. Gene enrichment analysis is performed to identify biological pathways, gene ontology terms, or other functional categories that are enriched in the set of genes of interest using clusterProfiler. The RNAseq data have been deposited to the Gene Expression Omnibus (GEO) platform and are available with the accession number, GSE284552.

#### Histology and immunofluorescence microscopy

Paraffin-embedded lung tissues were sectioned at 10-µm thick, stained with hematoxylin and eosin (H&E), Masson’s Trichrome (MT), or Immunohistochemistry (IHC) for SARS-CoV-2 nucleocapsid protein (Thermo Fisher, MA5-36251), and imaged on a Nikon Eclipse 80i, Nikon Plan Apochromat 20x objective lens (numerical aperture 0.75). For immunofluorescence microscopy, some lung tissues were inflation fixed for 12 hours with paraformaldehyde (4%, buffered neutral) at 4°C. Lungs were then placed in 30% sucrose for ≤1 week and embedded in optimal cutting temperature compound (Fisher Scientific, 23730571) in prelabeled Cryomold squares (VWR, 25608916) followed by dry ice for 10 to 15 minutes. After sectioning, slides were dried at room temperature for 1 to 2 hours before permeabilization, blocking, and immunostaining. Sections were incubated with α-tubulin Alexa Fluor 488 (5 μg/mL; Thermo Fisher, 322588), human ACE2 (5 μg/mL; R&D Systems, MAB10823) with Donkey anti-Rabbit IgG (H+L) Secondary Antibody, Alexa Fluor 594 (8 μg/mL; Fisher Scientific, A21207), CD68-APC (2μg/mL; BioLegend, 137008), CD115 Alexa Fluor 488 (5 μg/mL; BioLegend, 135511), or myTags SARS-CoV-2 Labeled for (-) Strand, 3X ATTO 550 (Daicel Arbor Biosciences, 451202), and then mounted with Hoechst 3342 (Biotium, 40046) and antifade mountant (Thermo Fisher, P36930). Using a Nikon Eclipse 80i inverted microscope with Nikon Plan Apochromat 10x objective lens (numerical aperture 0.3), 3-4 fluorescence images were acquired in the regions of interest as determined by cellular infiltration using the Hoechst staining. The area positive for hACE2 protein was quantitated in each airway field. The quantitation of cell types was expressed by the absolute count of each cellular marker (CD68, CD115, FISH, hACE2) per high-powered field or by the ratio of double positive cells to the total CD68^+^ or CD115^+^ stained cells in the field. Each individual data point represented one animal and reflected the average of 3-4 images captured from adjacent tissue sections per slide. All subsequent quantitative imaging analyses were performed using Image J software (112).

### QUANTIFICATION AND STATISTICAL ANALYSIS

All statistical data were analyzed using Prism 10 (GraphPad, v10.1.0). Results are presented as mean (SD) for continuous variables with normal distribution or equal variance and as median (interquartile range) for continuous variables without normal distribution. The *t*-test compared normally distributed continuous variables between 2 independent groups. Differences between multiple groups were assessed by ANOVA followed by Tukey’s post hoc multiple comparisons test. The Mann-Whitney *U* test was used for continuous variables not normally distributed. Male and female animals were used in equal numbers, and sex was analyzed as a biological variable, by 2way ANOVA.

