## Supplemental Figures for "IL-1β-driven NF-κB transcription of ACE2 as a Mechanism of Macrophage Infection by SARS-CoV-2"

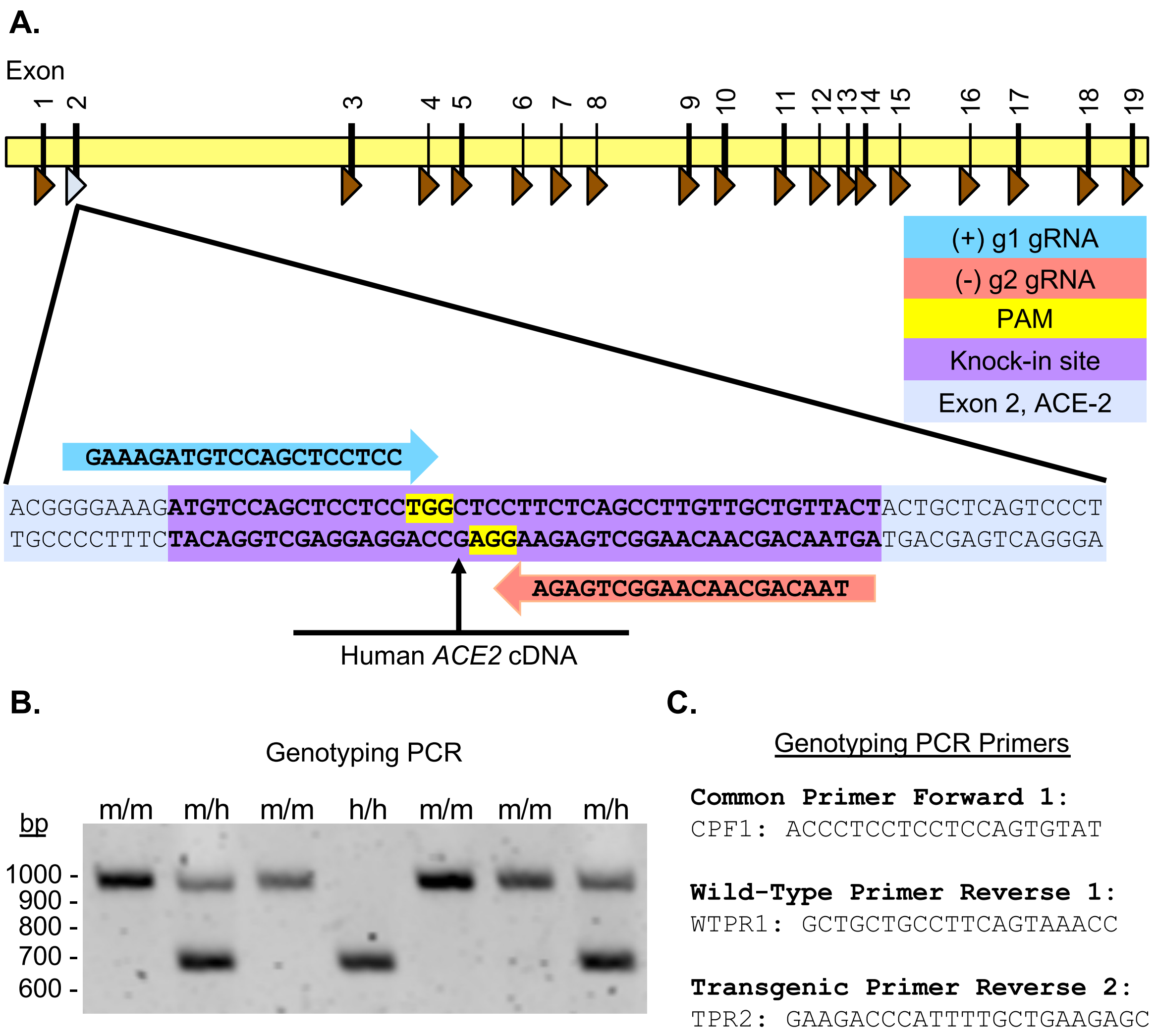


**Figure S1. Diagram of *hACE2* Gene Editing and Genotyping PCR identifying *hACE2* Mice.** (A) *hACE2* Gene Editing Strategy. Two recognized guide sites ((+) g1, light turquoise, (-) g2, rose) in exon 2 (light blue). Yellow shaded letters represent the protospacer adjacent motif (PAM) of 5'-NGG-3'. Purple represents the knock-in region. After the CRISPR/Cas9 system introduced a double strand break in two targeted sites of the *ACE2* allele, the homologous donor vector recombined, which resulted in the insertion of the human *ACE2* cDNA sequences. (B) Genotyping PCR products, using the common forward (CPF1) and wild-type reverse (WTPR1) and transgenic reverse (TPR2) primers respectively, were then resolved by electrophoresis on 2% agarose gel to differentiate mice with wild-type exon 2 from mice with human *ACE2* cDNA insertion. Expected PCR product size for mouse, wild-type allele (m) is 969 bp and for transgenic humanized *ACE2* insert allele (h) is 687 bp. (C) The sequences of the genotyping PCR primers.

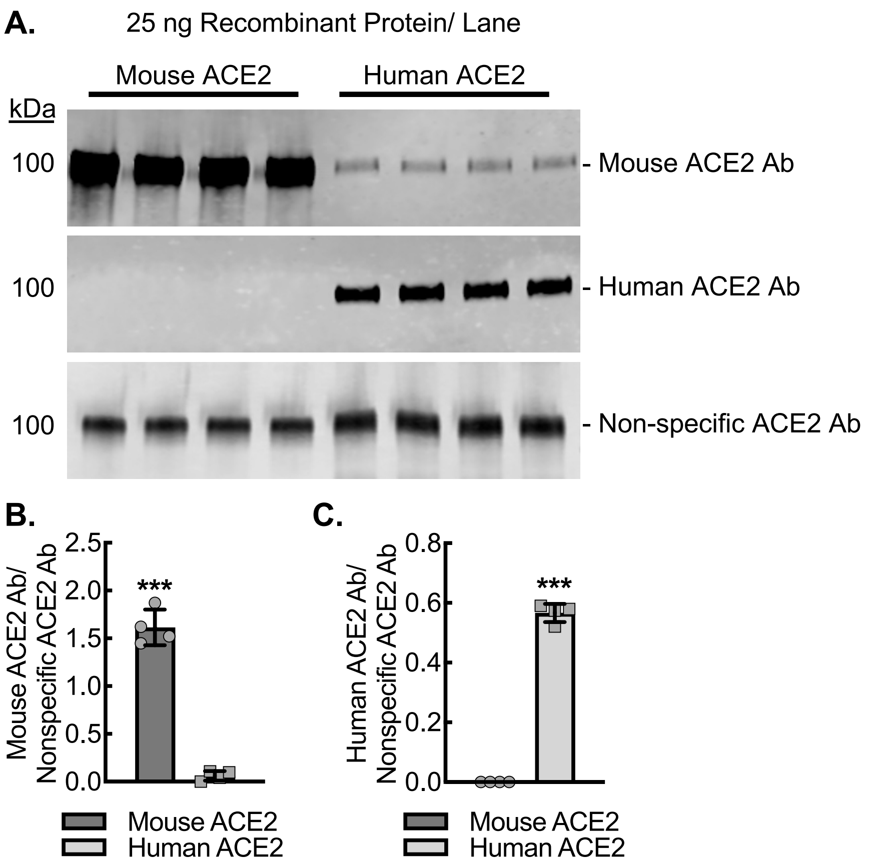


**Figure S2. Validation of antibodies specific to mouse or human ACE2 protein.** (A) Representative immunoblots loaded with 25 ng of recombinant mouse or human ACE2 per lane followed by staining with primary antibodies directed at mouse ACE2, human ACE2, or nonspecific primary antibody directed at ACE2 protein independent of species along with quantitative densitometry analysis (B, C). (***, P<0.0001; by t-test, n=4). Data, mean ± SD.

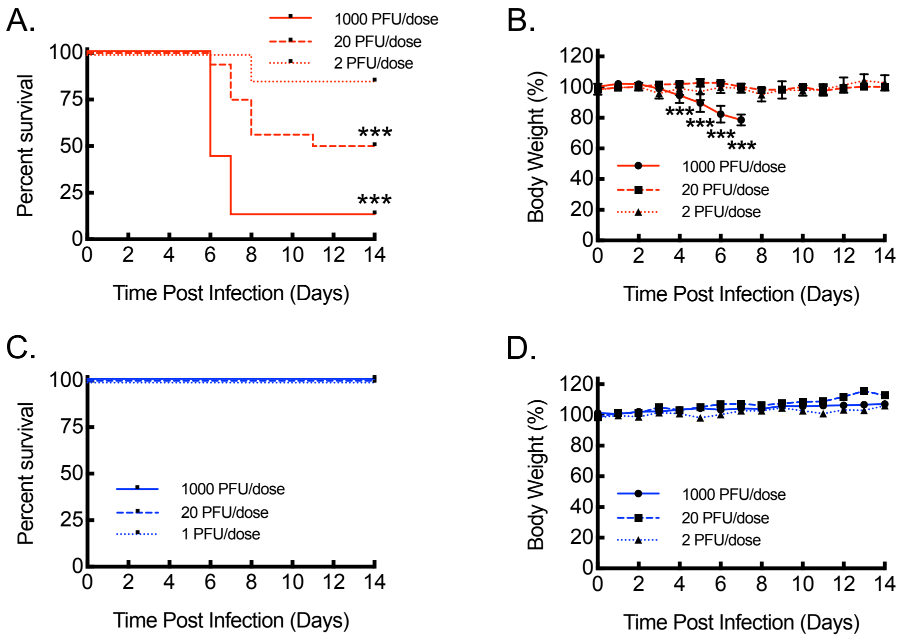


**Figure S3. Dose-Response Curves for Infection of *K18-hACE2* and *hACE2* mice with SARS-CoV-2**. (A) Kaplan-Meier plot for survival post intranasal infection of *K18-hACE2* mice (red) with SARS-CoV-2 at indicated concentrations of undiluted (1000 PFU/dose; n=16, 8 male and 8 female), 1:50 dilution (20 PFU/dose) (n=8, 4 male and 4 female), and 1:500 dilution (2 PFU/dose) (n=8, 4 male and 4 female) (***, P<0.0001 by log rank). (B) Percent body weight relative to pre-infection weight over time after SARS-CoV-2 infection carried out as in (A) (***, P<0.0001 by ANOVA). (C) Kaplan-Meier plot for survival post intranasal infection of *hACE2* mice (blue) with SARS-CoV-2 at indicated concentrations of undiluted (1000 PFU/dose; n=16, 8 male and 8 female), 1:50 dilution (20 PFU/dose) (n=8, 4 male and 4 female), and 1:500 dilution (2 PFU/dose) (n=8, 4 male and 4 female). (D) Percent body weight relative to pre-infection weight over time after SARS-CoV-2 infection carried out as in (C).

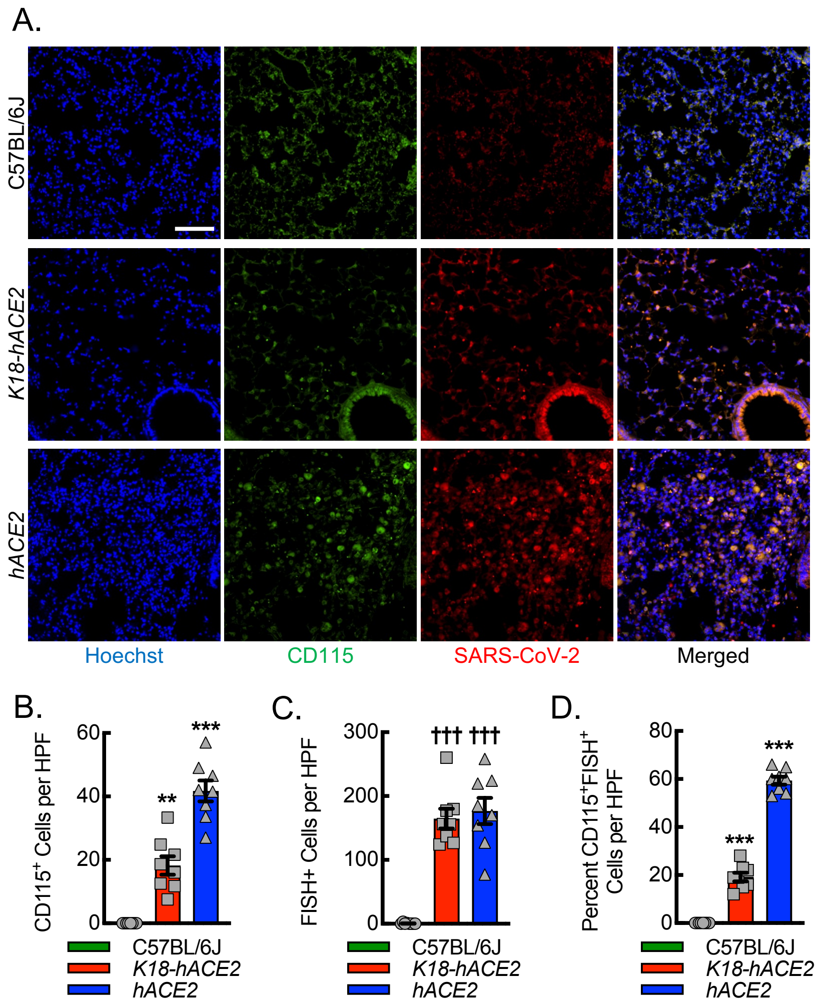


**Figure S4. SARS-CoV-2 replicates more efficiently in macrophages of hACE2 mice relative to K18-hACE2 mice.** (A) Representative immunofluorescent photomicrographs from lungs of C57BL/6J, *K18-hACE2*, and *hACE2* mice harvested on day 6 post intranasal infection with 1000 PFU/dose SARS-CoV-2. Blue, Hoechst. Red, FISH for SARS-CoV-2 negative strand. Green, CD115. Bar, 100 microns. Quantitation of the number of $CD115^{+}$ cells (B), number of $FISH^{+}$ cells (C), and the percent of $CD115^{+}$ cells that are also $FISH^{+}$ cells (D). (**, P<0.0005; ***, P<0.0001 compared to all others using ANOVA; †††, P<0.0001 compared to C57BL/6J using ANOVA; n=8, 4 male and 4 female).

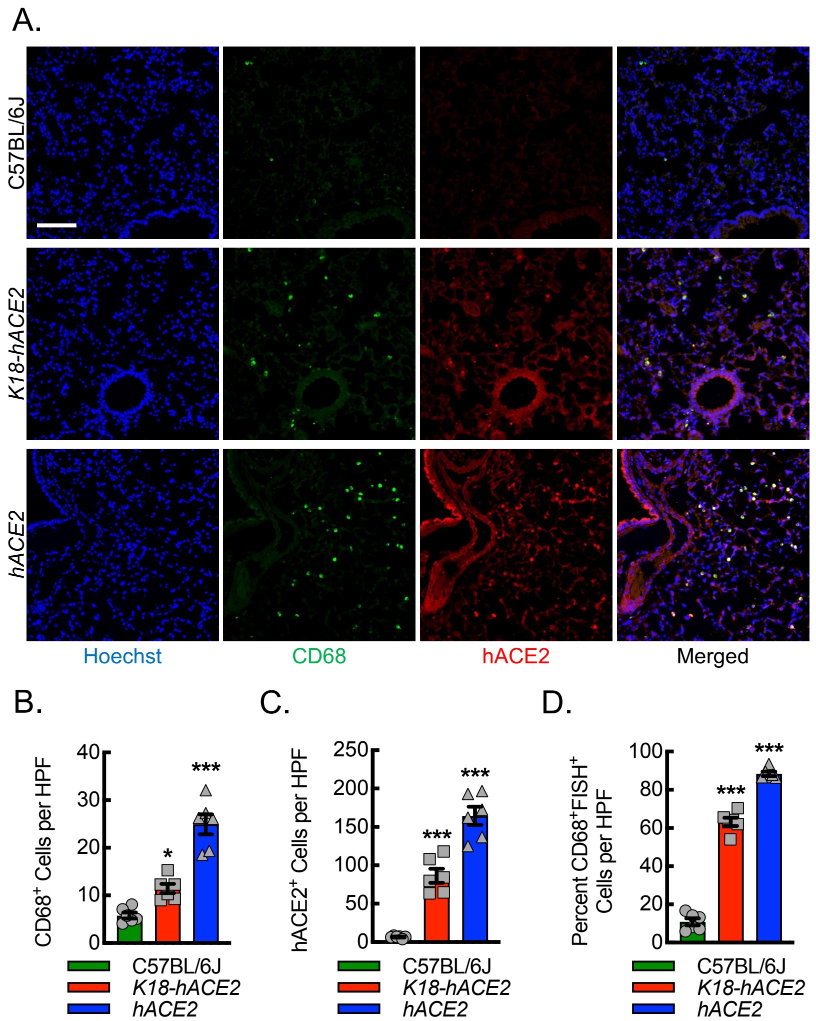


**Figure S5. CD68⁺ macrophages and the number of hACE2⁺ CD68⁺ macrophages are increased in lungs of hACE2 mice infected with SARS-CoV-2.** (A) Representative immunofluorescent photomicrographs from lungs of C57BL/6J, *K18-hACE2*, and *hACE2* mice harvested on day 6 post intranasal infection with 1000 PFU/dose SARS-CoV-2. Blue, Hoechst. Red, hACE2. Green, CD68. Bar, 100 microns. Quantitation of the number of $CD68^+$ cells (B), number of $hACE2^+$ cells (C), and the percent of $CD68^+$ cells that are also $hACE2^+$ (D). (*, P<0.05; ***, P<0.0001 compared to all others using ANOVA; n=6, 3 male and 3 female).

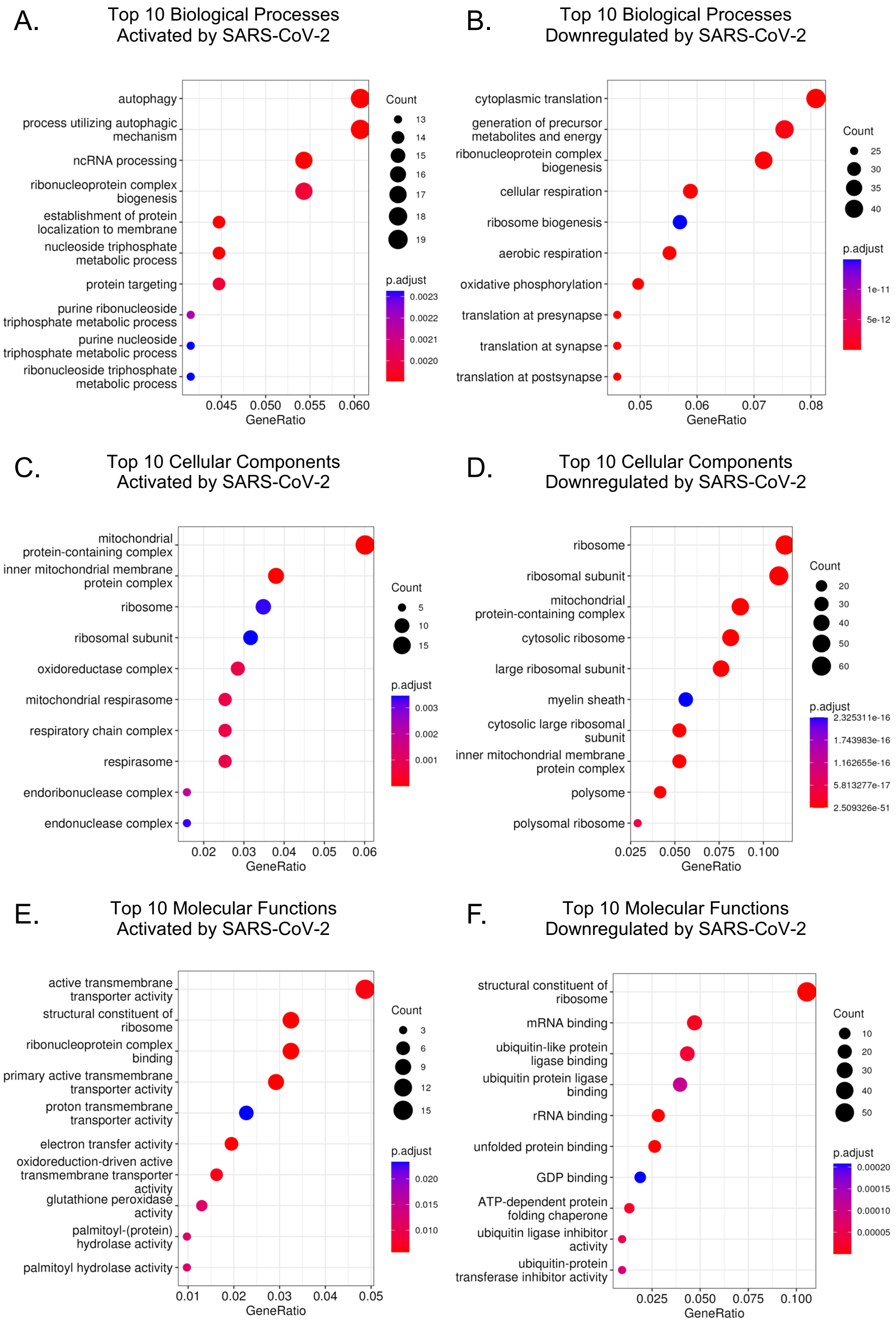


**Figure S6. Gene ontology analyses from differential transcriptomics of inflammatory macrophages infected with SARS-CoV-2 reveals alterations in RNA processing, protein translation, autophagy, mitochondrial function, and oxidation-reduction pathways.** Dot plots for the top 10 most significantly enriched upregulated (A, C, E) or downregulated (B, D, F) Gene Ontology terms derived from the differential transcriptomes between mock and SARS-CoV-2 infected BMDMs polarized with LPS+IFN-γ, comprising orthogonal ontologies of biological process (A, B), cellular component (C, D), or molecular function (E, F) (n=3 male mice) . Size, gene count. Color scale, adjusted P value.
